# Mapping Potential Antigenic Drift Sites (PADS) on SARS-CoV-2 Spike in Continuous Epitope-Paratope Space

**DOI:** 10.1101/2021.06.07.446560

**Authors:** Nathaniel L. Miller, Thomas Clark, Rahul Raman, Ram Sasisekharan

## Abstract

SARS-CoV-2 mutations with antigenic effects pose a risk to immunity developed through vaccination and natural infection. While vaccine updates for current variants of concern (VOCs) are underway, it is likewise important to prepare for further antigenic mutations as the virus navigates the heterogeneous global landscape of host immunity. Toward this end, a wealth of data and tools exist that can augment existing genetic surveillance of VOC evolution. In this study, we integrate published datasets describing genetic, structural, and functional constraints on mutation along with computational analyses of antibody-spike co-crystal structures to identify a set of potential antigenic drift sites (PADS) within the receptor binding domain (RBD) and N-terminal domain (NTD) of SARS-CoV-2 spike protein. Further, we project the PADS set into a continuous epitope-paratope space to facilitate interpretation of the degree to which newly observed mutations might be antigenically synergistic with existing VOC mutations, and this representation suggests that functionally convergent and synergistic antigenic mutations are accruing across VOC NTDs. The PADS set and synergy visualization serve as a reference as new mutations are detected on VOCs, enable proactive investigation of potentially synergistic mutations, and offer guidance to antibody and vaccine design efforts.

**Graphical Abstract:** 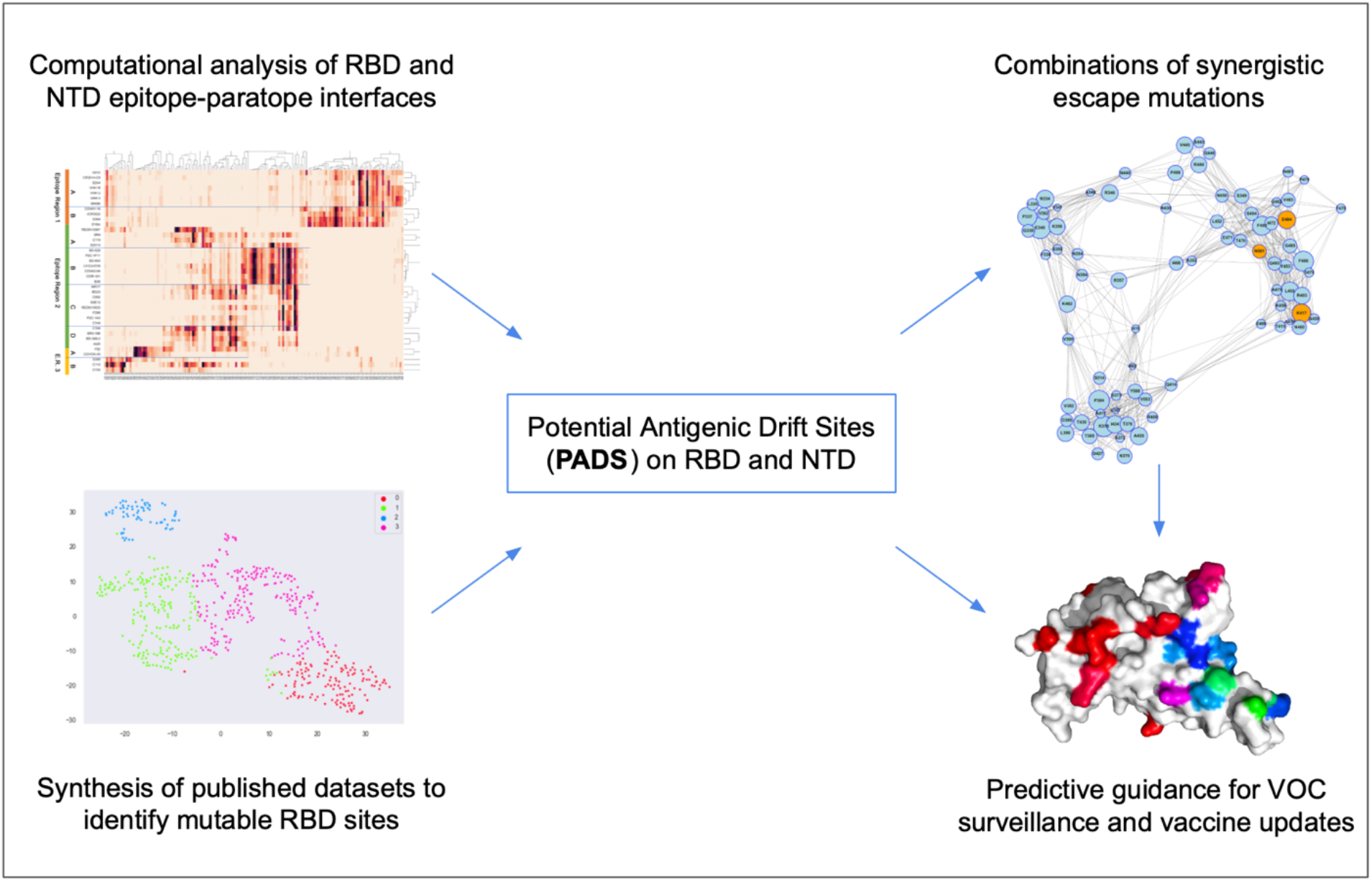

## Introduction

While an unprecedented sequencing effort to track SARS-CoV-2 (SARS2) evolution is underway, current surveillance efforts could be enhanced by additional experimental and computational tools. The identification of variants of concern (VOC) that are more transmissible [Davies et al., 2021], virulent [Challen et al., 2021], and partially resistant to antibodies [Wang et al., 2021] and immunity acquired through convalescence [Sabino et al., 2021] and vaccination [Madhi et al., 2021] emphasizes the critical need for surveillance. Though the VOC identified so far appear unlikely to severely infect convalescent individuals or to significantly jeopardize the current rollout of highly-effective vaccines [Abdool Karim et al., 2021], observation of convergent evolution of antigenic mutations on these VOC in combination with studies of seasonal coronaviruses [Eguia et al., 2021; Kistler et al., 2021] suggests that mutations will accumulate across the spike protein in response to selection pressure from host immune responses. A similar phenomenon has been reported in the case of influenza A viruses wherein mutations to escape host immune response are assimilated as the virus transmits between individuals who are antigenically naive and those who have been exposed via infection or vaccination [Cobey et al., 2017]. SARS-CoV-2 presents a unique opportunity to study mutations that could lead to antigenic drift over a much shorter timeframe than what is typically observed for influenza, which has been circulating in humans for decades. Although the currently observed VOCs might not represent true drift mutations that cause a substantial portion of people with exposure to the original antigen (through infection or vaccination) to become reinfected and have severe disease [Abu-Raddad et al., 2021], they do represent mutational paths for evolution of antigenic drift. Therefore, there is a need to develop tools to understand and potentially predict the evolving landscape of antigenic mutations in the receptor binding domain (RBD) and N-terminal domain (NTD) of the spike protein towards guiding vaccine development efforts and public health responses.

The RBD and NTD present numerous antibody epitopes that are highly overlapping and variably engaged, and it would be beneficial to derive a continuous description of “epitope-paratope space” at these domains to enhance our understanding of the key antigenic residues and the relationships between them. RBD epitopes have thus far been described using a variety of approaches, including the classification of antibodies targeting the RBD into four major “classes” based on structural analysis [Barnes et al., 2020]. This discrete description has proven useful for understanding key VOC mutations: the E484K mutation was associated with escape from class 2 antibodies while mutations at K417 were associated with escape from class 1 antibodies [Greaney et al., 2021b]. However, discrete descriptions do not convey heterogeneity within nor overlap between epitopes/mAb classes. For example, it is not clear to what extent there remains class 1 and 2 antibody pressure on VOCs bearing the E484K and K417N/T mutations, especially from antibodies that do not fit neatly into these classes or that partially overlap with another class. Similarly, it is difficult to ascertain the degree to which emerging mutations that provide escape from a specific group of antibodies such as L452R (which escapes binding of certain antibodies in class 3 but also has a slight effect on certain class 2 antibodies [Greaney et al., 2021b]), might provide synergistic escape with VOCs bearing E484K (class 2 escape). Therefore, there is a need for alternative continuous descriptions of RBD epitope-paratope space—as well as extension to NTD given the dominant role NTD Abs can play for some individuals [Voss et al., 2021]—to better understand and potentially predict the landscape of mutations that are antigenically synergistic with mutations on current VOCs.

In this study, we sought to map potential antigenic drift sites (PADS) using an epitope-paratope space representation that illustrates the potential for evolution of synergistic escape mutations in the RBD and NTD sites. Specifically, we aimed to build a model of PADS that could be used to understand the potential for newly observed mutations to provide synergistic escape with mutations on current VOCs, as such synergistic mutations are most likely to reduce the protective margin provided by convalescence and vaccination. Toward this end, we (1) performed a computational analysis of co-crystal structures of mAbs and nanobodies complexed with RBD and NTD to map epitope-paratope space at these domains; (2) defined a set of sites on RBD that are unconstrained to mutate based on genetic, structural, and functional features drawn from existing experimental datasets [Starr et al., 2020], GISAID [Shu et al., 2017], and our own structural analysis; (3) defined the set of PADS as those residues that are both antigenic and unconstrained to mutate; and (4) integrated (1), (2), and (3) to build a model of PADS in continuous epitope-paratope space that facilitates interpretation of potential synergistic escape between mutations on RBD and NTD. The resulting set of PADS provides a reference set of RBD and NTD sites of elevated antigenic drift risk, and the model of PADS offers a continuous map of RBD/NTD epitope-paratope space. While our results add immediate value toward understanding why certain mutations appear selected together (e.g., E484K and K417N/T) while others do not (e.g. E484K and L452R), our model will provide additional value as mutations accrue on VOCs and the scientific community seeks to understand the potential antigenic impact of each one. Further, the continuous epitope-paratope space model can aid proactive surveillance of VOCs, for example, via assessing whether convalescent, vaccinee, and VOC booster sera neutralizes pseudoviruses bearing VOC + predicted synergy mutations, as well as serve as the basis for immune-focusing vaccine design.

## Results

### Epitope-Paratope Analysis of RBD-mAb Structures Captures Binding Escape from Convalescent Sera

Toward our first goal of mapping RBD and NTD epitope-paratope space from co-crystal structures, we leveraged and extended a previously validated computational approach for quantifying protein networks, known as Significant Interaction Networks (SIN) [Soundararajan et al., 2011]. Briefly, mAb-RBD/NTD co-crystal structures are converted to network-models, with nodes describing residues and edges describing non-bonded interactions between residues. From the network model of epitope-paratope complexes, we derive networking scores between all pairs of epitope and paratope residues. While a direct networking metric has been described and validated previously for measuring interactions spanning the epitope-paratope interface [Robinson et al., 2015], we theorized that an indirect networking metric would better capture allosteric effects critical to assessing the impact of a given epitope mutation on antibody escape (see *Methods*). To benchmark our indirect networking metric, we measured the correlation between indirect networking (computed across mutable RBD residues for 39 RBD-mAb complexes) and a published experimental dataset describing binding escape from convalescent sera [Greaney et al., 2021a] (Figure S1). We found a strong correlation (r=0.65, p < 0.01) and observed that indirect networking captured all mutable residues with elevated sera escape, suggesting that the epitope-paratope analysis utilized in this study is largely representative of the polyclonal sera response measured in Greaney et al. 2021a.

### RBD and NTD Epitope-Paratope Mapping

Next, we applied SIN to the set of mAb/nanobody-RBD/NTD complexes to produce networking scores between each mAb/nanobody and every residue on RBD (Figure 1) and NTD (Figure S2). Direct and indirect networking scores were normalized mAb-wise and summed to obtain a total networking score between each mAb and each RBD/NTD residue, and the resulting epitope-paratope networking matrices were clustered epitope-wise (RBD/NTD residues) and paratope-wise (mAbs and nanobodies). For RBD, the mAb clustering (Figure 1, X-axis dendrogram) indicates three “epitopes regions” (ERs) on RBD, with each ER containing 2-4 overlapping epitopes (labeled A-D for each ER). For NTD, the mAb clustering indicates a single dominant epitope region that is consistent with previous descriptions of the NTD “supersite” [Cerruti et al., 2021], as well as a second under-sampled NTD ER that is so far only structurally-characterized for a single mAb (DH1052, [Li D et al. 2021]).

**Figure 1:**
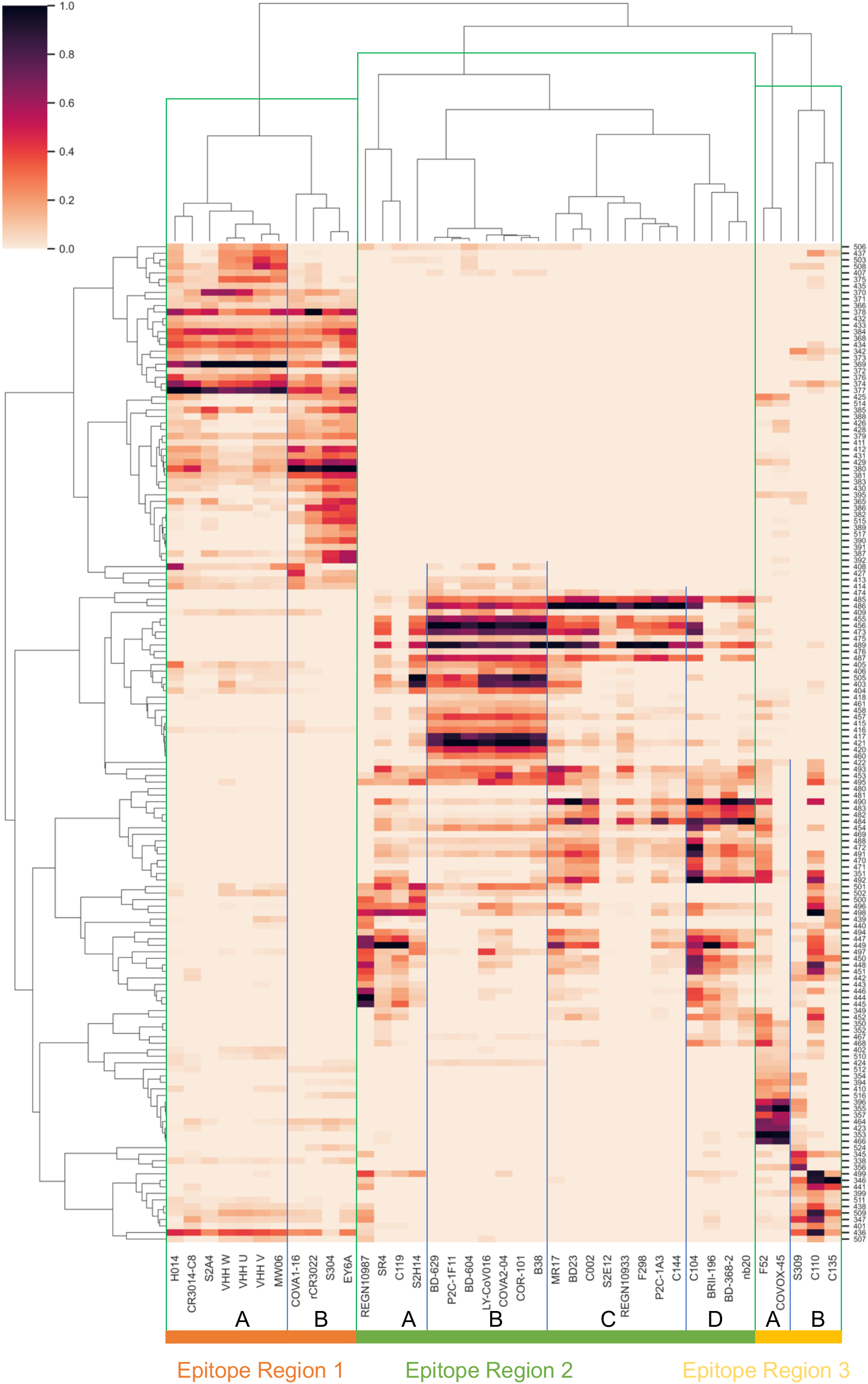
Computational RBD Epitope-Paratope Mapping from mAb- and Nanobody- RBD Complexes. The color of each square represents total networking between a given mAb paratope (x-axis) and RBD residue (y-axis). To aid visualization, only residues that score in the top 75% of all residues for at least three of the mAbs or nanobodies are shown. Equivalent maps for direct and indirect networking in isolation are included in the supplement (Fig S1-2). Three epitope regions (ERs 1-3) are highlighted as orange, green, or yellow and are separated by vertical green lines. Two to four epitopes (A-D) are annotated within each ER and separated by vertical blue lines, reflecting groups of highly similar mAbs and nanobodies. While certain ERs are highly uniform (ER1-B), others are highly variable (ER2-C).

The three RBD ERs appear to broadly correspond to the RBD back (ER1), ACE-2 binding site (ER2), and RBD front (ER3). The epitopes within each ER significantly overlap yet also indicate substantial differences which reflect, among other features: distinct epitope residues, relative networking strength of key residues, and variations in mAb/nanobody angle of attack. ER1 broadly describes the SARS-conserved epitope region on the RBD back that is only accessible in RBD up and further consists of two overlapping yet distinct epitopes. Interestingly, despite significant overlap, these two epitopes appear to neutralize via different mechanisms, with antibodies targeting epitope ER1-A neutralizing via avidity [Liu et al., 2020] and those targeting ER1-B neutralizing via ACE2 binding interference [Lv et al., 2020]. ER2 describes the immunodominant ACE2-binding site and includes both the most mAbs and the most epitopes. Antibodies targeting these epitopes overwhelmingly bind both RBD up and RBD down and neutralize primarily via steric obstruction of ACE2 binding. Meanwhile, ER3 describes the relatively poorly characterized RBD epitopes on the RBD front that are distinct from the ACE2 binding site. Our map divides ER3 into two sub-epitopes, but the low number of mAbs and high degree of heterogeneity suggest that this ER remains under-sampled in the Protein Data Bank (PDB). While this may be largely explained by the fact that mAbs targeting this region such as COVOX-45 tend to be weak neutralizers [Dejnirattisai et al., 2021], others such as S309 potently neutralize [Pinto et al., 2020]. The epitope-paratope map highlights how COVOX-45 and S309 differentially interact with ER3 RBD residues, offering insights into the ER3 residues enabling potent neutralization for S309.

In contrast to ER3, the high number of structurally-characterized mAbs and nanobodies in ER2 (ACE2-binding surface) likely results from the selection bias during mAb isolation and development for the most potent neutralizers, which tend to target the ACE2 binding site [Jiang et al., 2020]. A similar phenomenon appears to exist for NTD with all but a single (weakly binding) NTD mAb in PDB targeting the same “supersite” ER. Still, substantial variation across mAbs targeting RBD ER2 exists allowing identification of approximately four distinct epitopes within ER2: ER2-A, ER2-B, ER2-C, and ER2-D. In particular, mAbs targeting ER2-B feature the least amount of epitope-paratope networking variation likely reflecting evolutionary proximity to the common germline IGHV3–53 [Huang et al., 2020; Raybould et al., 2020], while the other epitopes within ER2 feature greater variation. Similar to the S309 and COVOX-45 comparison above, the epitope-paratope map facilitates straightforward comparison of mAbs targeting similar or overlapping RBD and NTD epitopes, which can be correlated with experimental readouts to generate mechanistic hypotheses for differences in affinity, pan-coronavirus breadth, escape-susceptibility, or other features.

Despite the overrepresentation of mAbs targeting the ACE2 binding site (ER2), the epitope-paratope map still indicates coverage of mAbs targeting the other two ERs. Thus, in combination with our observation that indirect networking captures the major RBD escape mutations observed in Greaney et al. 2021a, we hypothesized that the epitope-paratope map covers the dominant antibody epitopes on RBD and NTD and can serve as the basis for 1) identifying potential antigenic drift sites (PADS) within these domains, and 2) defining a continuous representation of epitope-paratope space for estimation of synergy between PADS.

### Computation of Potential Antigenic Drift Sites (PADS)

After mapping RBD and NTD epitope-paratope interactions, we next sought to determine the set of PADS before finally projecting the PADS into continuous epitope-paratope space for interpretation of synergistic mutations. We defined PADS as those mutations which we predict to have 1) weak constraints on mutation, and 2) an antigenically relevant mutation effect. While a lack of mutational constraints is a requirement for mutation, our approach is not predicated on nor predictive of any specific selective pressure such as host immune response or fitness enhancement. Rather, we sought to identify residues that are not genetically, structurally, or functionally constrained to mutate. We integrated computational analyses, existing experimental datasets, and log mutation frequencies in GISAID to estimate genetic, structural, and functional constraints on RBD site mutation. Note functional mutagenesis datasets are so far not available for NTD as they are for RBD, and NTD appears tolerant of substantial mutation and deletion based on the variety of NTD mutations observed on VOC. Consequently, identification of NTD PADS was based entirely on antigenicity analysis.

For RBD PADS, we first applied a single nucleotide polymorphism (SNP) filter to remove all RBD amino acid mutations that are not achievable via a SNP from the reference SARS-CoV-2 sequence (Wuhan-Hu-1). Next, we applied spectral clustering (see *Methods*) to identify higher-dimensional relationships across the computational and experimental mutability features (Figure S5, Table S1). The set of RBD SNP mutations was well described by four clusters, one of which (Cluster 0) is dominantly mutable, and three of which are primarily immutable according to the average characteristics of the residues in each cluster. We filtered out the constrained mutations belonging to the primarily immutable clusters, taking the unconstrained mutations belonging to the dominantly mutable cluster forward to our antigenic analysis. (Figure S5, Cluster 0).

Finally, we sought to identify residues that have the greatest antigenic effect upon mutation within the set of unconstrained and low-fitness cost mutations. To estimate antigenic effect, we calculated a single antigenicity score as the weighted sum of three epitope-paratope interface features (Figure S5, Table S2), where the weights for each feature are the coefficients of the first principal component of the feature matrix. Two of these features are previously introduced as direct and indirect networking and describe the importance of a given residue to epitope-paratope interfaces and within-epitope structure. To estimate the magnitude of perturbation by mutation to a specific residue, we added a third feature representing the change in epitope-paratope surface complementarity (SC) upon mutation (see *Methods*). Importantly, all three features are computed using the set of mAb/nanobody-RBD complexes which contains a significant PDB sampling bias in the number of mAbs per ER, suggesting normalization may be required to compare the antigenicity of residues from different ERs. On the other hand, as previously mentioned our analysis suggests this bias is roughly correlated with neutralization potency due to the large number of structurally characterized mAbs targeting the ACE2 binding site. However, it is precisely these epitopes that have been knocked down by VOC mutations such as E484K and K417N/T, and so we believe the most accurate contemporary depiction of RBD and NTD epitope-paratope space is one in which these biases are controlled for, such that less immunodominant sites that may become increasingly relevant on future VOC are highlighted. Therefore, we correct for the PDB sampling bias by normalizing the antigenicity for each RBD/NTD residue to the number of mAbs directly or indirectly networked to the given residue. This workflow generates normalized antigenicity estimates for each of 79 unconstrained RBD residues—the PADS set. We next interpret the PADS in continuous epitope-paratope space to build our understanding of potential antigenic drift paths for the SARS-CoV-2 RBD and NTD.

### Network Model of PADS in Continuous Epitope Space

Having identified the PADS set, we sought to project the PADS into a continuous epitope-paratope space that encompasses the direct and indirect networking between residues in the RBD and NTD and the entire set of mAbs and nanobodies, allowing identification of key PADS and the relationships between them. Briefly, we converted each residue in the set of PADS to a node in our network and defined node positions and edges according to distance between residues in epitope-paratope space. Distances between pairs of residues in epitope-paratope space are close for residues that bind highly similar sets of mAbs with similar relative strengths and distant for residues that bind orthogonal sets of mAbs (see *Methods*). For an illustrative example, consider residues E484 and F490. As shown in Figure 1, the set of mAbs that are networked to E484 are nearly the same set of mAbs networked to F490. Further, E484 and F490 score similarly high for most of the mAbs they interact with relative to other residues. Therefore, mutations at F490 are unlikely to provide synergistic escape with mutations at E484 since mutations at both residues likely affect a highly overlapping set of mAbs, and F490 should be closely associated with E484 in epitope-paratope space. On the other hand, escape mutations at residues which bind entirely orthogonal sets of mAbs would be likely to synergistically knock down binding on VOCs harboring the E484K mutation, and thus will be distant from one another in the network representation of epitope-paratope space. This workflow produces a network model of PADS in epitope-paratope space that enables interpretation of the potential for synergistic escape between residues based on relative node positions. We present both the full network model for the entire set of PADS (Figure S6), as well as a reduced network model for just the top 50% most antigenic PADS on RBD and the top 75% most antigenic PADS on NTD (Figure 2).

**Figure 2:**
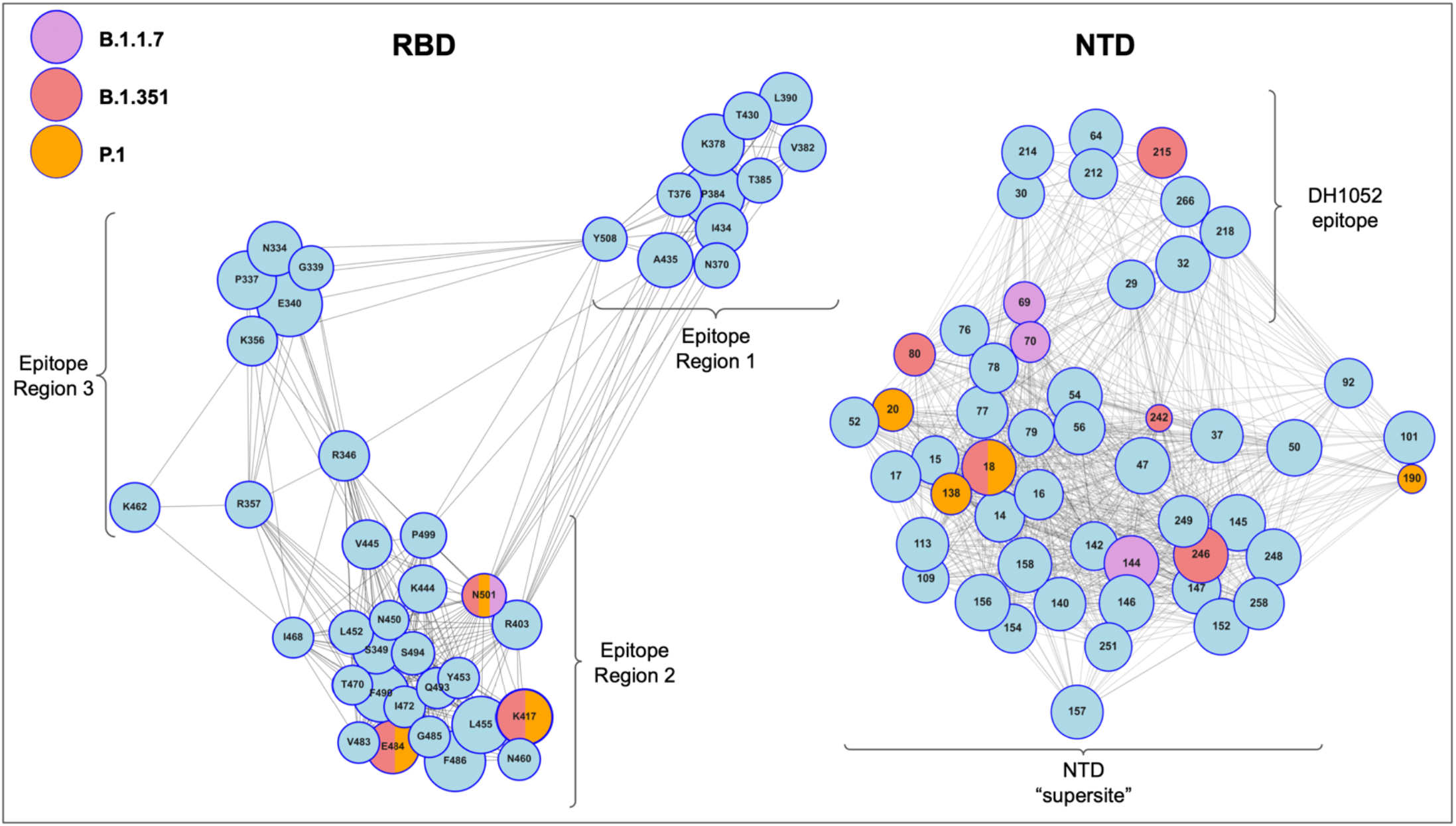
RBD and NTD PADs in Continuous Epitope-Paratope Space. Each RBD or NTD PAD is represented as a node, with node size proportional to estimated antigenicity. Node positions and edges are determined based on cosine similarity between sites in epitope-paratope space (Figure 1). Only the PADS scoring in the top 50% of antigenicity for RBD and 75% for NTD are shown for interpretability (all PADS shown in Figure S5). The three main clusters RBD residues map to the three RBD ERs shown in Figure 1. The NTD clusters map to the NTD ‘supersite’ and the DH1052 epitope. Centrally-networked PADS are key across the ER they are contained within (e.g., E484), while PADS residing on the ER fringes describe residues that are uniquely important to certain epitopes or mAb binding modes within that ER (e.g., K417). Key centrally-networked RBD sites are identified in ER1 (K378, P384) and ER3 (P337, E340), as well as unique sites that that span ERs (R346). While VOCs exhibit *identically* convergent evolution at RBD sites (K417, E484, N501), NTD mutations appear *topologically* convergent in epitope-paratope space (e.g., 142/144/246 and 20/80).

The network model of RBD PADS broadly includes the epitope regions defined in Figures 1 and S2 and enables further analysis of within ER and between ER epitope relationships, such as escape synergy potentials between sets of mutations. Residues that are centrally-networked within epitope-paratope space clusters appear to be fundamental across most mAb epitopes within the corresponding epitope region, meaning that mutations at these residues have the potential to disrupt a large subset of the mAbs targeting this epitope-region. We see evidence for this effect in residues E484, T470, I472, and S494 which are centrally-networked within the PADS groups representing ER2, consistent with experiments demonstrating knockdown of mAbs with distinct binding modes upon mutation [Zhou et al., 2021; Rees-Spear et al., 2021]. In contrast, residues that are on the fringes of epitope-paratope space for a given ER cluster appear uniquely important to a specific epitope or mode of mAb binding within that ER which logically builds escape synergy between mutations at these sites and the more centrally-networked PADS within these groups. For example, the triplet K417, E484, and N501 geometrically spans most of ER2 (Figure 2) suggesting that each mutation is playing a minimally-overlapping role in disturbing mAbs targeting ER2. Likewise for NTD, B.1.351 mutations L18F, D80A, and R246I span much of the supersite while D215 is centrally networked within the distinct DH1052 epitope. That is, VOC mutations appear efficiently spread out across epitope-paratope space suggesting that antigenic synergy likely plays a role in the selection of these mutations and that the epitope-paratope space model accurately represents this synergy.

### The Epitope-Paratope Space Network Model Highlights Synergy Between Key VOC Escape Mutations

We next interpret the network model by examining the key RBD VOC mutations at K417, E484K, and N501. Mutations at residue E484 and K417 occur on the VOCs that are most strongly associated with escape from vaccinated individuals (501Y.V2; [Madhi et al., 2021]), convalescent individuals (P.1; [Sabino et al., 2021]), and vaccine sera and mAbs [Wang et al., 2021; Zhou et al., 2021]. Similarly, N501Y is found on nearly all VOCs and is more strongly associated with enhanced ACE-2 binding than antigenic escape, though a moderate escape effect is noted for a subset of mAbs [Zhou et al., 2021]. These hypotheses align well with our measure of antigenicity at these sites, in which the node size (estimated antigenicity) of K417 and E484 stand out strongly within the network and are greater than the node size of N501. Our analysis thus supports hypotheses that mutations at N501 are driven primarily by factors other than immune evasion, as they score relatively low on antigenicity but that N501 may still contribute somewhat to escape. Further, the network indicates that K417 is closely associated with E484 in global RBD epitope-paratope space (within the ER2 cluster), but relatively distant from E484 within ER2 (*cos(θ) =* 0.13; Figure 3A). This finding is consistent with recent reports classifying K417N/T and E484K as knocking down different discrete antibody classes [Greaney et al., 2021b]. Further, K417 is relatively isolated from other RBD residues within ER2 suggesting that K417 interacts uniquely with a subset of mAbs within ER2.

**Figure 3:**
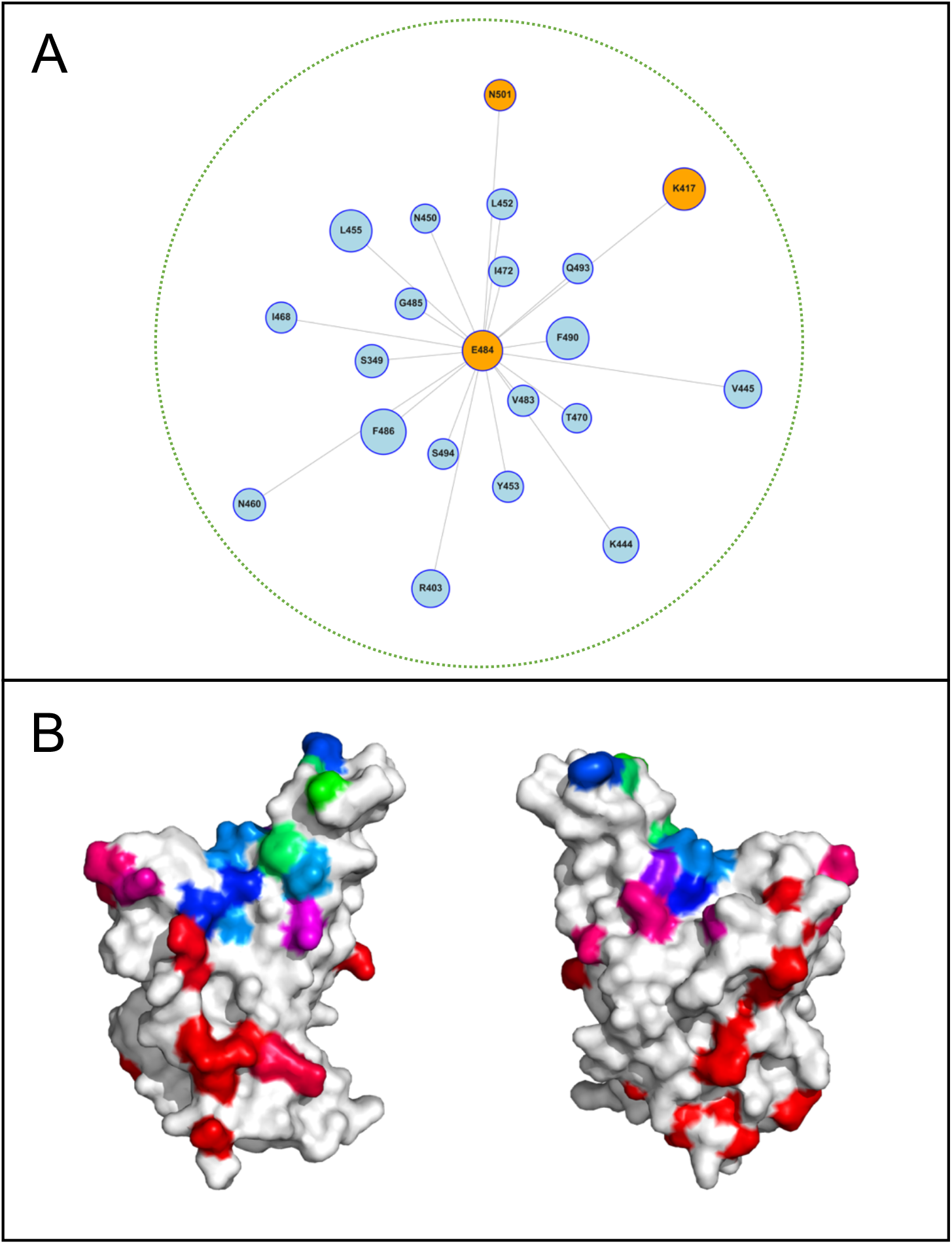
Most Antigenic PADS According to Epitope Distance from E484. (A) A sub-model containing only the ER2 PADS is shown with edges and node positions determined solely based on epitope distance from E484. This model isolation simplifies interpretation of potential synergy between mutations at residues within the ACE2 binding site and E484. Notably, within the E484-adjacent group K417 and N501 are relatively distant from E484. L452 is relatively closer to E484 yet still further than immediately adjacent sites such as F490 and F486. (B) The PADS with the top 50% of antigenicity scores are shown on RBD, and colored according to their distance from E484 in epitope space. Colors map as follows: PADS that are E484-overlapping in epitope space are highlighted in green (networked to a highly similar set of mAbs), E484-proximal in blue (networked to a partially overlapping set of mAbs), E484-intermediate in magenta, and E484-distant in red (networked to a distinct set of mAbs).

L452R was identified as a defining mutation on B.1.617 alongside E484Q [Cherian et al., 2021] as well as on other VOC including B.1.427 and B.1.429. Our analysis identifies L452 as antigenic and moderately associated with E484 (*cos(θ) =* 0.53, Figure 3A) compared to immediately adjacent residues such as E484/F490 (*cos(θ) =* 0.90) and more distant residues such as E484/K417 (*cos(θ) =* 0.13). This suggests that mutations at L452/E484 have the potential for escape synergy but significantly less so than E484/ K417 and E484/N501. A limitation of our approach is the difficulty in defining a similarity cutoff that denotes clinically-relevant synergy. Based on experimental evidence and convergent evolution, low similarities approaching those of K417/E484 (0.13) are very likely synergistic and a cutoff may eventually prove inferable via observation of VOC evolution. It is interesting to note that L452R has to our knowledge only been observed alongside E484Q and not in combination with E484K. Indeed, E484K appears to disrupt mAb binding to a more significant degree than E484Q across multiple readouts, including our surface complementarity analysis (data not shown), published escape mapping experiments [Greaney et al., 2021a], and examination of chemical properties (EàK vs. EàQ). It is plausible, then, that L452R is relatively more synergistic with E484Q than it is with E484K. If this hypothesis is supported by additional observations of L452R alongside E484Q (but not E484K), then a similarity cutoff slightly above 0.53 may be appropriate for RBD ER2 as sites sharing this degree of mAb overlap could require specific mutational conditions for relevant synergy. Note also that the mutation T478K, observed on B.1.617.2, occupies a highly unique fringe position in epitope-paratope space (Figure S6), but is not one of the highest scoring PADS included in Figure 2. Still, other mutations within ER2 appear to have a clearer basis for synergy with E484.

### Future Synergistic Mutations Within or Proximal to the ACE2 Binding Surface and NTD Supersite

In addition to K417 and N501, residues on the 443-450 loop also occupy a relatively distant position within the E484-proximal group (ER2) with highest antigenicity estimated for mutations at K444 (*cos(θ) =* 0.17) and V445 (*cos(θ) =* 0.11). Residues on this loop have been shown to have high binding escape from sera and have been proposed as future escape mutants on VOCs [Greaney et al., 2021a]. Our antigenicity and synergy analyses are consistent with these predictions given that these sites measure similarly to K417 and N501 in terms of distance from E484 in epitope-paratope space. With E484/(444, 445) cosine similarities lower than E484/K417, we expect these sites to be sufficiently orthogonal to provide escape synergy in combination with mutations at E484. Further, these sites are also distant from K417 (*cos(θ) =* 0.06), indicating that mutations at K444 and V445 may prove synergistic with VOCs bearing both E484K and K417N/T. Whether these two sites on the 443-450 loop or others which we predict to have lesser antigenicity and lesser synergy such as 450 are mutated will also depend on fitness considerations not examined here beyond our determination that these sites are not substantially constrained to mutate—a limitation of this work. However, we note that certain sites on this loop such as N450 are substantially closer to E484 in epitope-paratope space (*cos(θ) =* 0.52). Other PADS within or adjacent to ER2 that appear distant in our network model from both E484 and K417 include N439, S443, G446, P499, and I468/R346, though the last two are somewhat associated with ER3.

Examining NTD, we find a potential explanation for the experimentally observed differences between VOCs B1.1.7, B.1.351, and P.1. As mentioned above, the B.1.351 mutations at sites 18, 80, 215, and 246 appear to span the NTD supersite while also potentially knocking down the DH1052 epitope which contributes to protection *in vivo* despite a lack of *in vitro* neutralization [Li D et al., 2021]. In contrast, P.1 NTD mutations at sites 20 and 138 are concentrated around the more antigenic mutation at L18 suggesting a degree of antigenic redundancy for these mutations. The P.1 mutation at site 190 is distant from this triplet cluster, but residue 190 itself appears poorly antigenic and not well networked to many other antigenic residues (only closely associated with site 101). Indeed, these findings are consistent with comparisons of sera escape for P.1 and B.1.351 in which B.1.351 is measured to have higher sera escape despite the two VOCs sharing nearly identical RBD mutations (N501Y, E484K, and K417N/T; [Hoffmann et al., 2021]). Meanwhile, the B.1.1.7 NTD deletion at site 144 is highly central to the NTD supersite, consistent with reports of B.1.1.7 NTD escape. Further, deletion at sites 69-70 exist on the supersite edge that is most closely networked to the DH1052 epitope and it is plausible that the 69-70 deletion provides knockdown of antibodies targeting both the supersite and the distinct DH1052 epitope, similar to how R346 appears to span RBD ERs 1 and 2. While identically convergent evolution has not been observed for NTD as it has for RBD, our epitope-paratope space mapping provides evidence of *functionally* convergent NTD antigenic evolution. That is, in epitope-paratope space, both B.1.1.7 and B.1.351 share a similar topological constellation of mutations (144 mapping to 246, and 69-70 mapping to 18, 80, and 242), while P.1 so far lacks this constellation, and this fact may explain P.1’s slightly weaker escape from neutralization. Further, multiple B.1.617 lineages have emerged bearing a number of mutations within the NTD that are consistent with the proposed convergent evolution in epitope-paratope space. In particular, mutations at site 142 occupy a highly overlapping position with those observed at 144 and 246 on for B.1.1.7 and B.1.351. Similarly, mutations at site 19 are consistent with the mutations observed at sites 18 and 20 on for B.1.351 and P.1. In contrast, mutations and deletions at sites 154, 156, 157, and 158 occur on the southern fringe of the NTD supersite, a location in space not perturbed for other VOC. Still, these sites are predicted to be antigenic, indicating that B1.617 lineages may have enhanced escape relative to other VOC from mAbs targeting the NTD via the mode of binding mediated by this fringe, which is most strongly associated with CDRL3 of mAbs 4-18 and S2X333.

### Future Synergistic Mutations Distant from the ACE2 Binding Surface

The epitope-paratope space model highlights regions of epitope-paratope space which might provide synergistic escape with existing VOC mutations. There are two primary RBD PADS groups distal from E484, K417, and N501; one on the left-side of the network which is somewhat associated with the 443-450 loop and maps to ER3, and one in the upper right portion of the network which is largely isolated and maps to ER1. Antibodies binding residues where these mutations occur have essentially zero connection to E484/K417/N501 in epitope-paratope space and thus are most likely unaffected by mutations at these sites, unless via long-range allosteric interactions which are not measured by our indirect epitope networking calculation. Cosine similarity for all of these sites versus E484 is zero or nearly zero. Most of these sites reside on distinct RBD surfaces from E484 yet distance in epitope-paratope space does not map directly to surface distance, particularly within ERs (Figure 3B). While these sites have essentially zero connection to E484, a number of these sites have some degree of association with other PADS within ER2 (Figure S6).

Mutations at these VOC-distant PADS that prove to have an immune evasion effect are most likely to result in completely synergistic escape with variants bearing E484K and K417N/T. However, a potential “saving grace” is that a subset of these residues is not exposed in RBD-down and may be associated with escape from antibody binding but not neutralization [Greaney et al. 2021a]. Still, clinically-developed mAbs targeting these residues are capable of neutralization via avidity [Liu et al., 2020] or other mechanisms, and potently neutralizing antibodies targeting these residues have been isolated from the sera of convalescent individuals (EY1A, [Zhou et al., 2020]). Further, SARS-CoV-2 antibodies that fail to neutralize *in vitro* can still contribute to *in vivo* protection as evidenced by DH1052 [Li D et al., 2021]. A number of the residues in these groups such as P337, E340, and R346 are surface exposed and have been documented as escape mutants from neutralizing mAbs [Cathcart et al., 2021; Weisblum et al., 2020]. Antigenic mutations at these epitopes may only be significantly selected for after certain conditions are met, such as 1) dominant ACE2 epitopes have been sufficiently knocked down, and 2) a critical level of the population has been infected or vaccinated such that antigenic mutations provides a sufficiently high fitness advantage. Alternatively, novel epitopes formed by mutations at the ACE2 binding site may also prove more immunologically-relevant than these distal epitopes with relatively lower neutralizing potential, such that antibodies targeting the mutated ACE2 epitopes still dominate the sera response. Still, the identified sites at these epitopes are valuable to surveil. The distant residues that our model predicts to be the most antigenic and synergistic with VOC mutations are: N334, P337, E340, R346, K356, K378, P384, L390, I434, A435, V445, and K462, with lower ranking hits identified in the supplement (Figure S6). Similarly, for NTD our model suggests that P.1 might achieve additional escape from NTD antibodies by acquiring mutations at sites proximal to 144 and 246 in epitope space—for example sites 47, 142-147, and 246-248—as well as central to the DH1052 epitope (e.g. 212, 215, or 266).

### Mutations at PADS Have Been Observed on VOCs

VOCs have now been observed with mutations at a number of PADS. According to NextStrain as of 3/22/21, the 501Y.V2 lineage has mutations at PADS 348, 352, 359, 367, **382**, **384**, 408, 427, **430**, **434**, **435**, 439, 440, 443, **444**, **445**, 456, **450**, 457, 476, and 478, where bolded residues score in the top 50% of PADS [nexstrain.org]. In particular, P384 has 4 mutations to S/L on 501Y.V2 (*Ireland/D-NVRL-21IRL42037/2021, GISAID EPI ISL: 1117762, Zoe Yandle et al.; SouthAfrica/Tygerberg-446/2020, GISAID EPI ISL: 745179, Engelbrecht et al.; SA/NHLS-UCT-GS-9117/2020, GISAID EPI ISL: 1040685, Iranzadeh et al*). 501Y.V3 has few available sequences (375 in NextStrain as of 3/22/21), and no observed mutations yet at PADS. Many 501Y.V1 sequences are available, with mutations observed at PADS: **337**, 354, **356**, 367, 372, **376**, **390**, 394, 408, **430**, 440, **445**, 446, **455**, **468**, 471, 475, 478, **483**, **490**, and **494**. The epitope-paratope space model highlights how mutations observed on 501Y.V1—which does not have E484K—differ substantially from those observed on 501Y.V2. That is, 501Y.V1 lacks E484K and we observe mutations at sites that are in close proximity to E484 in epitope-paratope space (483, 490, 494) and so may be playing a similar antigenic role in the absence of E484 mutations. On the other hand, 501Y.V2 bears E484K and we have not yet observed mutations at proximal sites potentially due to antigenic redundancy with E484K.

## Conclusions

In this study, we have classified all RBD residues according to their role as continuous epitope constituents on RBD using mAb/nanobody-RBD/NTD complexes to approximate the polyclonal sera response. Further, we classified RBD mutations according to their estimated mutational constraints and scored mutable residues according to their estimated antigenicity to identify a set of potential antigenic drift sites (PADS) on RBD and NTD. Finally, we developed a continuous epitope-paratope space representation of the RBD and NTD PADS allowing visualization of each residue’s position in this space relative to all other RBD and NTD PADS. In particular, this model facilitates interpretation of the degree to which newly observed mutations might be antigenically synergistic with existing VOC mutations. The model provides evidence of topologically convergent evolution of NTD mutations across current VOCs. Further, the model assists interpretation and helps build intuition toward why certain VOC mutations are observed together such as E484K and K417N/T while others are not. We highlight PADS that offer the potential for synergistic escape with existing VOCs, both within and distant from the ACE2 binding surface and the NTD supersite. We advise careful observation of these sites on VOCs and highlight the potential for using these data to guide proactive surveillance and vaccine design.

## Methods

### RBD-mAb Networking Calculations

For each protein or protein-complex, significant interaction network (SIN) scores between every pair of residues within the structure were computed based on side-chain interactions as described previously [Soundararajan et al., 2011]. Given that the SIN computation is based on side chains, glycine networking was interpolated from nearby residues. Using the SIN output, the following metrics were defined. <u>Within RBD Networking</u>: The sum of all interactions between a given residue on RBD and all other residue on RBD. <u>Direct Paratope Networking</u>: For a given residue on RBD, the sum of all interactions between the RBD residue and all residues on the complexed antibody/nanobody. <u>Indirect Paratope Networking</u>: For a given residue on RBD, the sum of all interactions between the RBD residue and all other residues on RBD which are directly networked to an antibody/nanobody paratope. <u>Total Paratope Networking</u>: For a given residue on RBD, the sum of direct and indirect paratope networking scores. Note that for all networking measures, scores are normalized to the highest networking score within each RBD-mAb complex before they are subsequently compared. For subsequent computations that sought to control for mAb-epitope bias, networking scores at each epitope residue were normalized to the number of mAbs interacting with the given residue.

### RBD Mutations and Surface Complementarity Calculations

We used PyRosetta to model the impact of RBD SNPs on the surface complementarity of the RBD-mAb complexes [Chaudhury et al., 2010]. For each mutation, the wild type complex was first repacked, the mutation was performed, side chains were repacked, and the interface surface complementarity was computed using the Interface Analyzer Mover. Absolute surface complementarity perturbations were scored relative to the wild type interface surface complementarity.

### External Dataset Import and Processing

GISAID sequences were accessed and downloaded on 1/29/21 [Shu et al., 2017]. Mutation frequencies at all RBD residue were computed for every observed mutation. For ACE2 Binding [Starr et al., 2020], RBD Expression [Starr et al., 2020], and Sera Escape [Greaney et al., 2021a] datasets, raw data were downloaded from the Bloom Lab Github: https://github.com/jbloomlab. For all 3 datasets, the average values across both experiments and all patient samples were used for each mutation. For residue-level computations, mutation scores were averaged for SNPs only. The set of RBD SNPs was determined using a custom script and computed on the SARS-CoV-2 reference genome (NCBI RefSeq NC_045512.2).

### mAb-epitope clustering and Heatmap

<u>mAb-epitope clustering</u>: Direct, Indirect, and Total networking scores for each residue across all mAb and nanobody complexes were plotted using clustermap from the Seaborn statistical data visualization package with the ‘correlation’ distance metric and the ‘average’ linkage method [Waskom 2021]. <u>Mutability Clustering</u>: Mutability clustering of all RBD SNPs was performed using the computational and experimental features described in Table S1. GISAID mutation frequencies were log-transformed, all features were standardized, and then spectral clustering was performed on the affinity matrix generated using the pairwise Euclidean distances of the mutations based on the features (sklearn SpectralClustering [Pedregosa et al., 2011]). Clustering graphs were visualized using tSNE, and the accompanying cluster descriptions were computed as average scores for each feature for each cluster, except for the GISAID feature which was reported as the percentage of the cluster with an observed mutation in GISAID.

### Epitope Normalization and Epitope-Paratope Distance

As a preprocessing step starting from the raw networking matrix (Figure 1), we first apply the epitope bias correction, in which the networking scores for each residue are normalized using the number of mAbs and nanobodies interacting with each RBD/NTD residue. This step removes bias due to over- or under-sampling of mAbs/nanobodies interacting with a given epitope within our set of complexes. Next, we perform a residue-wise (row-wise) cosine similarity calculation between all pairs of residues. This process yields a single distance metric between all pairs of RBD residues, describing the distance between the pair in epitope-paratope space.

### Network Model for Antigenicity in Epitope Space

The most antigenic RBD mutation at each RBD site was chosen, and the top 50% of the most antigenic PADS are depicted in figure 2, while all PADS are shown in figure S6. Similarly, the top 75% of most antigenic NTD PADS were chosen. VOC mutants were included regardless of PADS status or antigenicity to highlight their location in epitope-paratope space. A network model was constructed in which node size is proportional to antigenicity from PCA (RBD) or total networking (NTD). Pairwise distance between residue-pairs in epitope-paratope space (see above method) was used to define edges and edge weightings, and was computed via the sklearn pairwise_distances function using cosine similarity as the distance metric [Pedregosa et al., 2011]. For clarity, edges are only draw between nodes with cosine similarity greater than 0.05 for the top PADS (Figure 2) and 0.25 for the full set of PADS (Figure S6). Node positions were computed via the Fruchterman-Reingold force-directed algorithm, which computes node position based on edge weights.

## Acknowledgements

NLM was supported in part by T32 ES007020/ES/NIEHS NIH.

## Author contributions

All authors conceived and designed the research. NLM and TC performed the research. NLM wrote the manuscript; all authors revised and approved the manuscript.

## Competing interests

The authors declare no conflicts of interest.

## Supplemental Figures (S1-S7) and Tables (S1 to S3)

**Figure S1:**
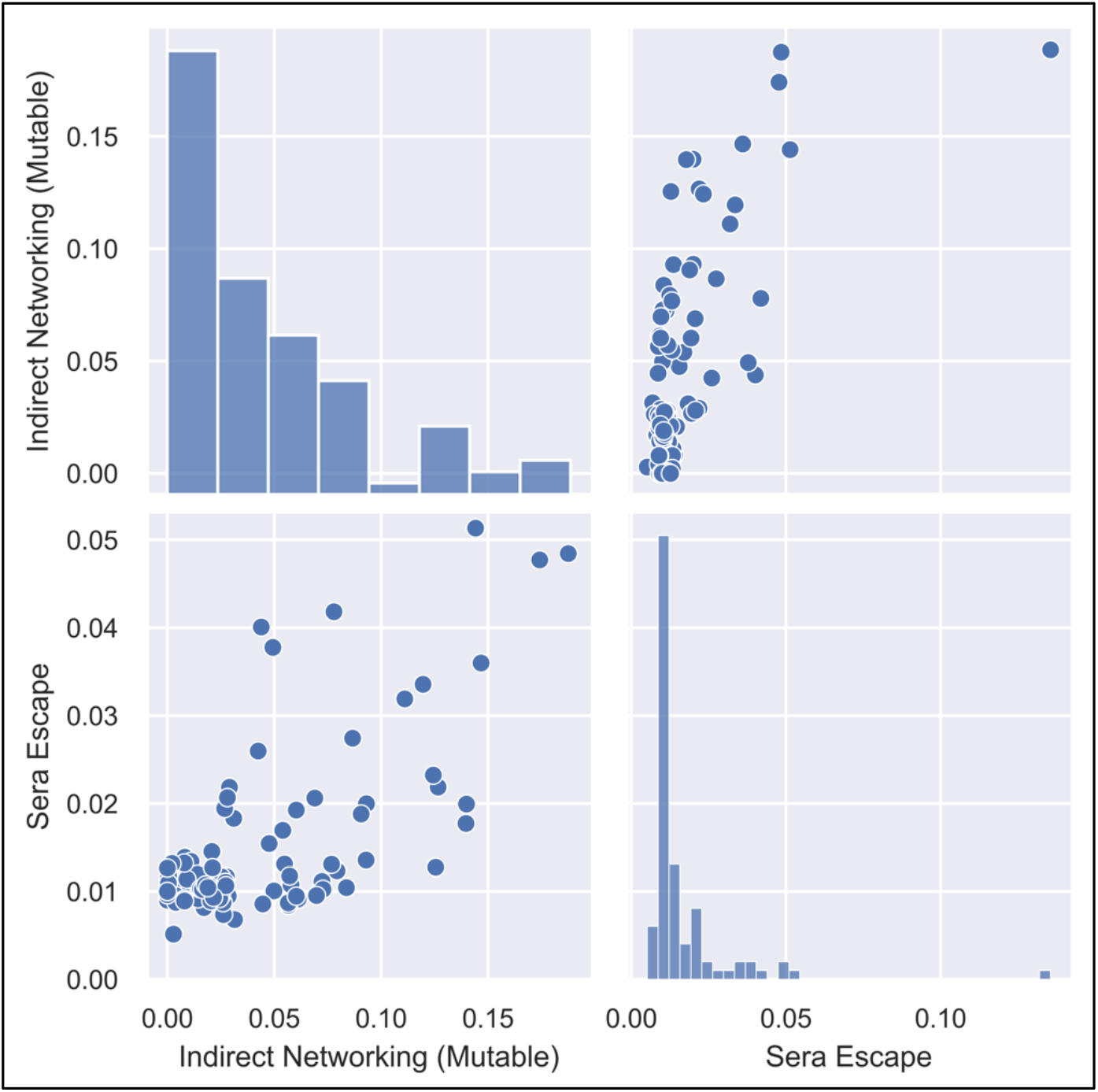
Correlation Between Indirect Networking of Mutable RBD Residues and RBD Sera Escape Mapping. Pearson correlation coefficient between sera escape [Greaney et al., 2021a] and indirect networking between mutable residues and paratopes of mAbs and nanobodies is r=0.65 (p < .01). For clarity, one outlying residue (E484) is removed from the bottom left graph, with approximate coordinates [0.19,0.14]. This point is shown on the top right graph, and is the highest scoring point for both sera escape and indirect networking.

**Figure S2:**
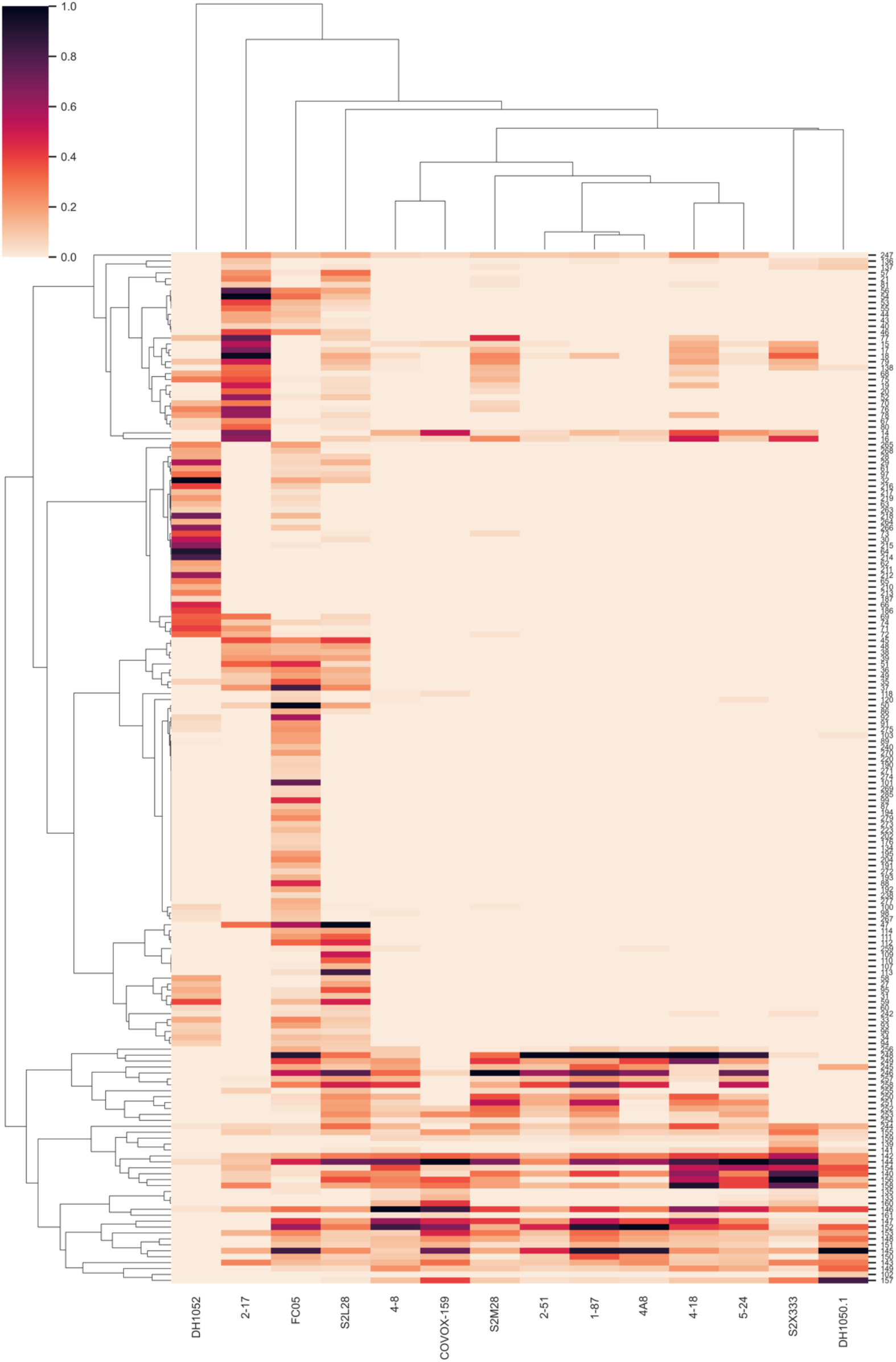
Total Epitope-Paratope Networking for N-Terminal Domain

**Figure S3:**
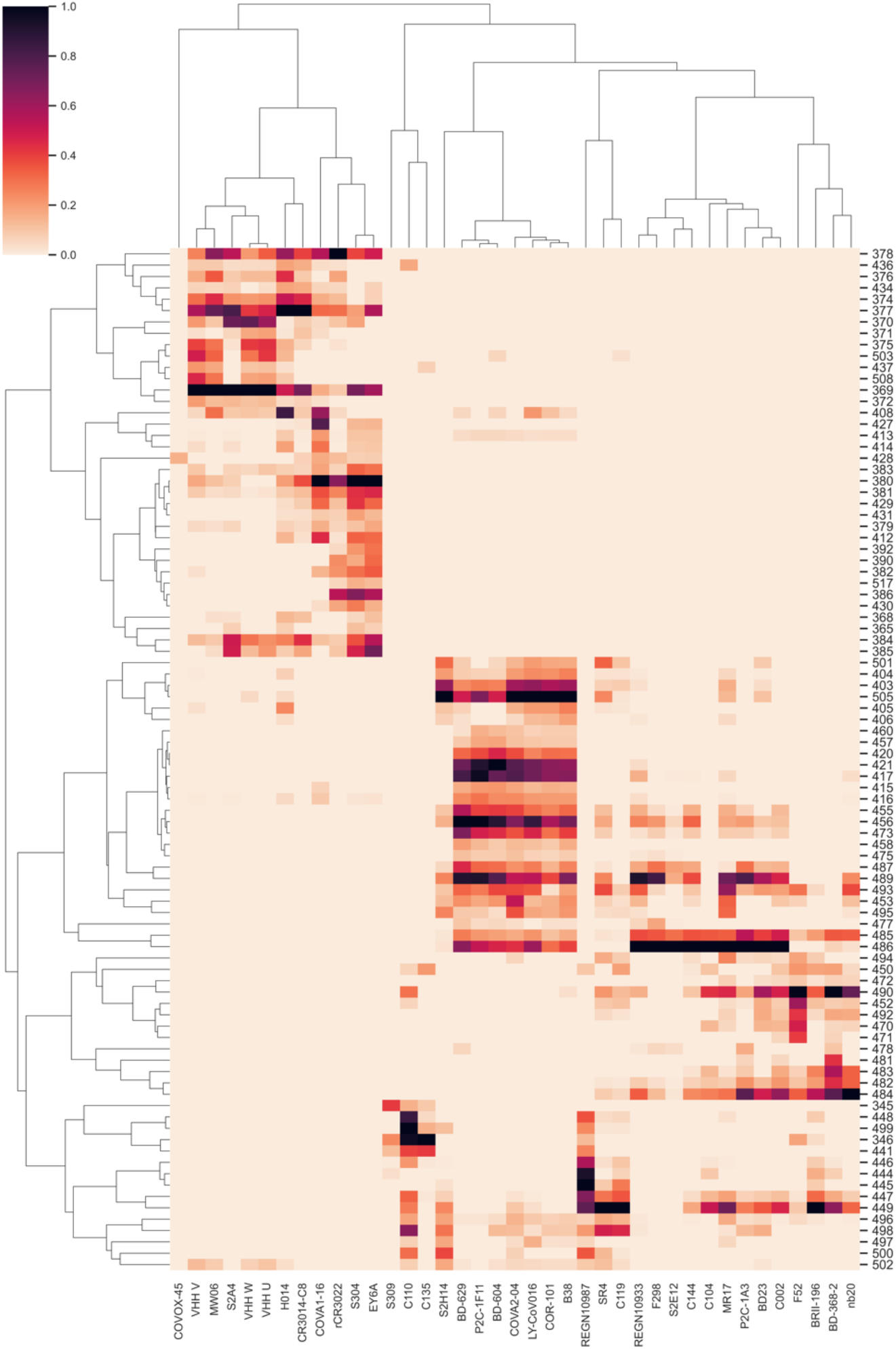
Direct Epitope-Paratope Networking for RBD

**Figure S4:**
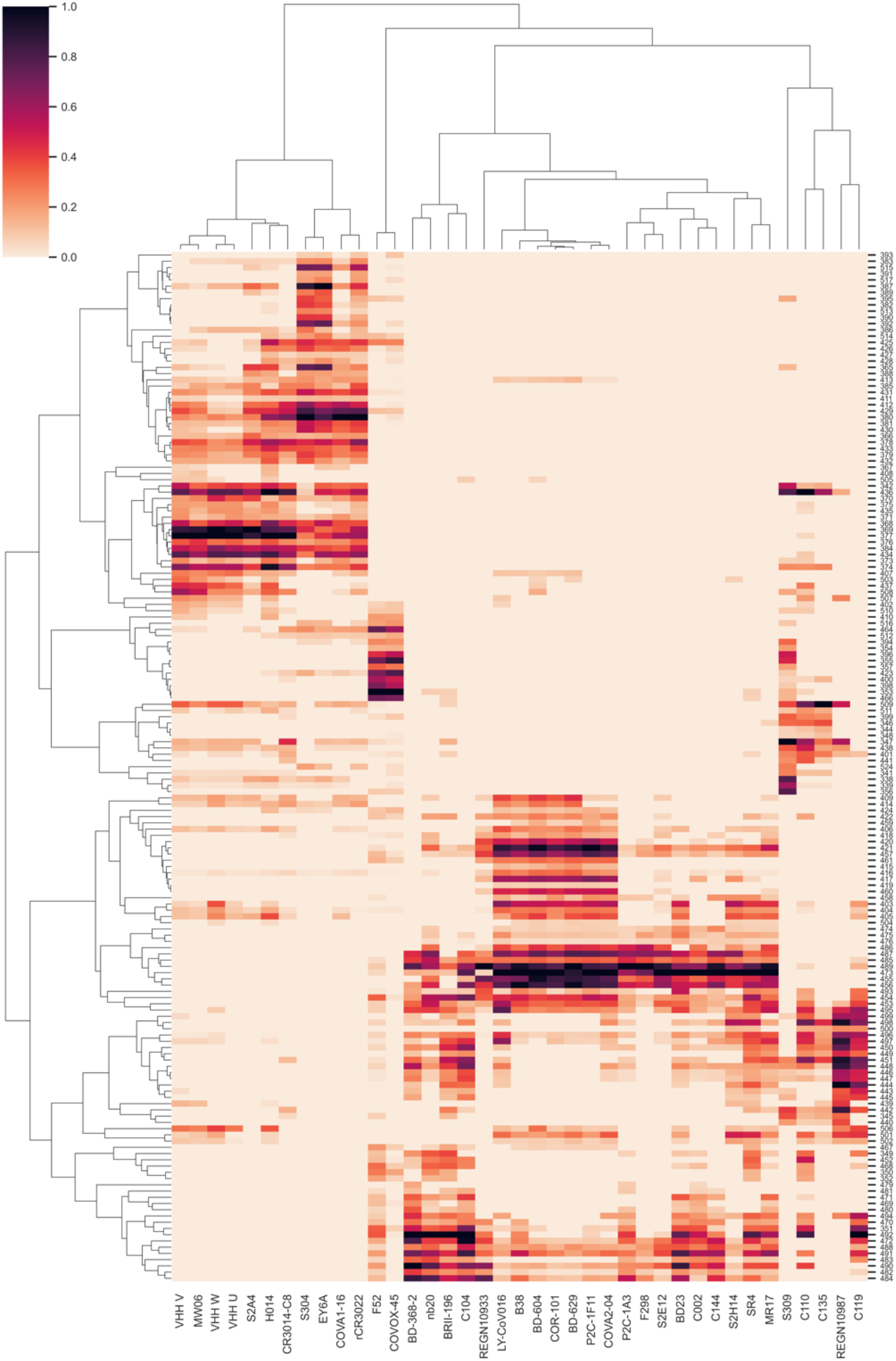
Indirect Epitope-Paratope Networking for RBD

**Figure S5:**
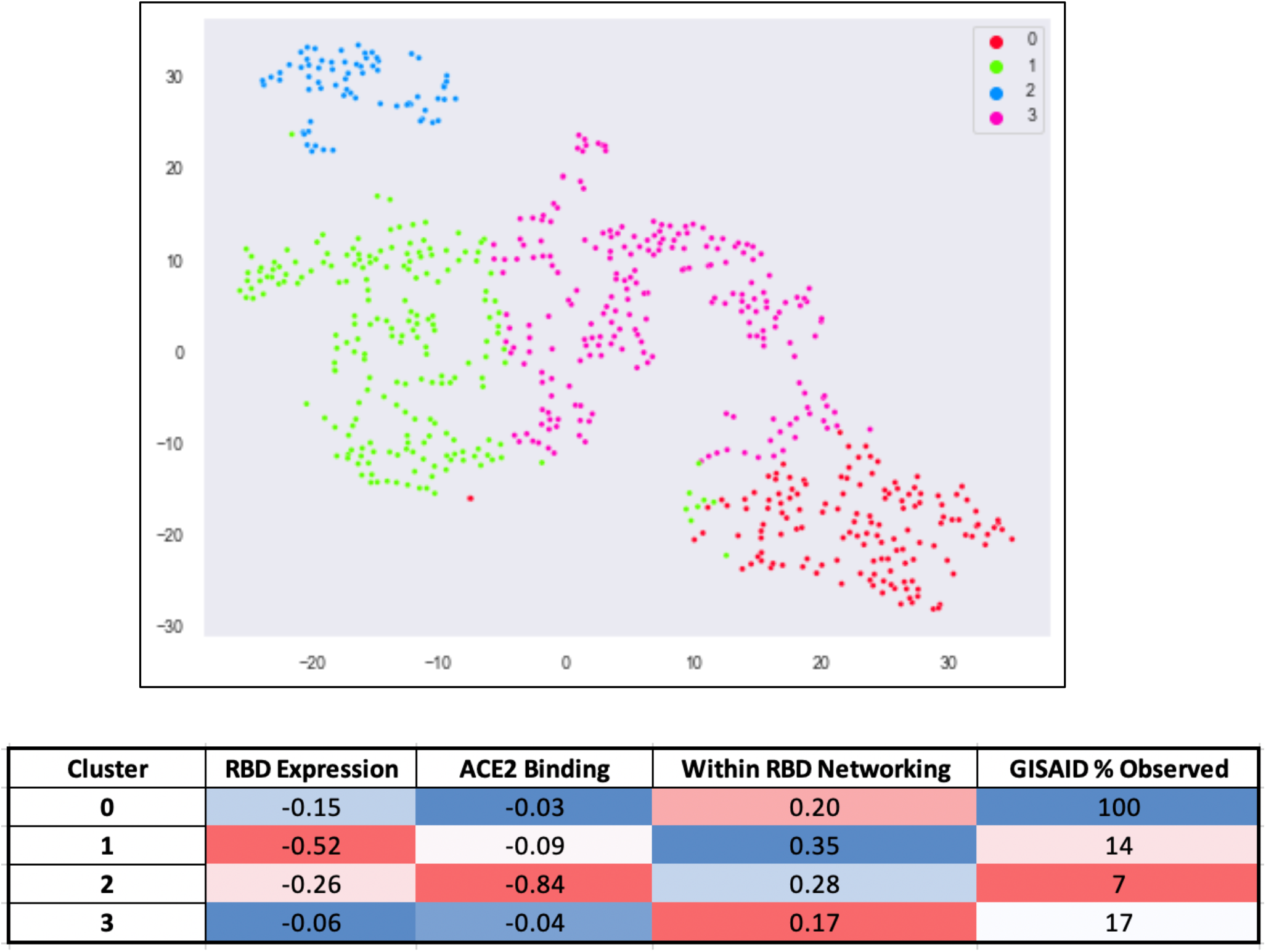
Mutability Clustering for all RBD SNP Mutations. Top, tSNE visualization of the clustering. Bottom, average and normalized (to absolute 1) values for each cluster along the feature set. Negative values imply knockdowns (e.g., in RBD expression, ACE2 binding, or ACE2 surface complementarity (SC)). Cluster 0 includes the RBD residues that appear the least genetically, structurally, and functionally constrained to mutate.

**Figure S6:**
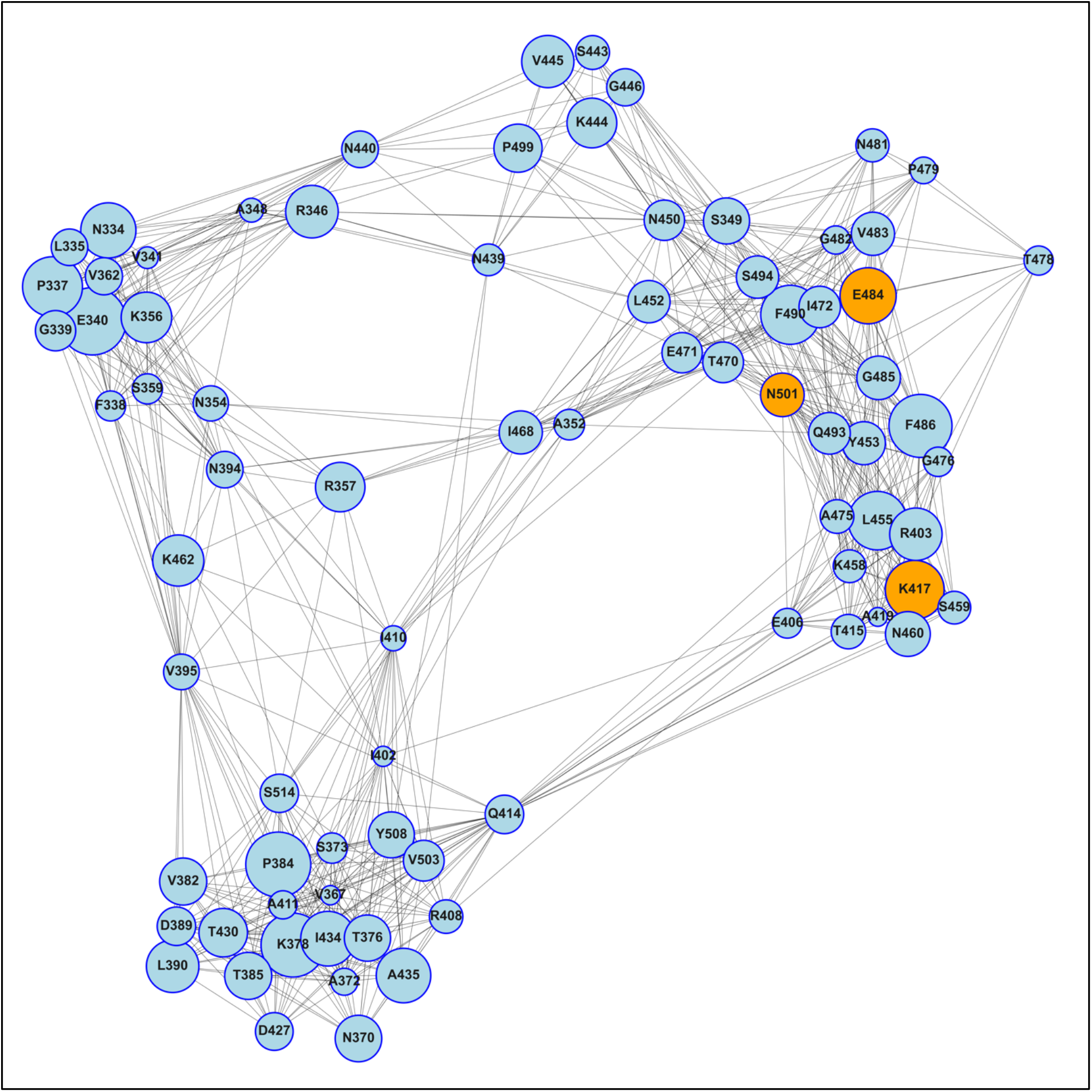
Network model depicting the entire set of RBD PADS, instead of only the top PADS shown in Figure 2. See Figure 2 legend.

**Figure S7:**
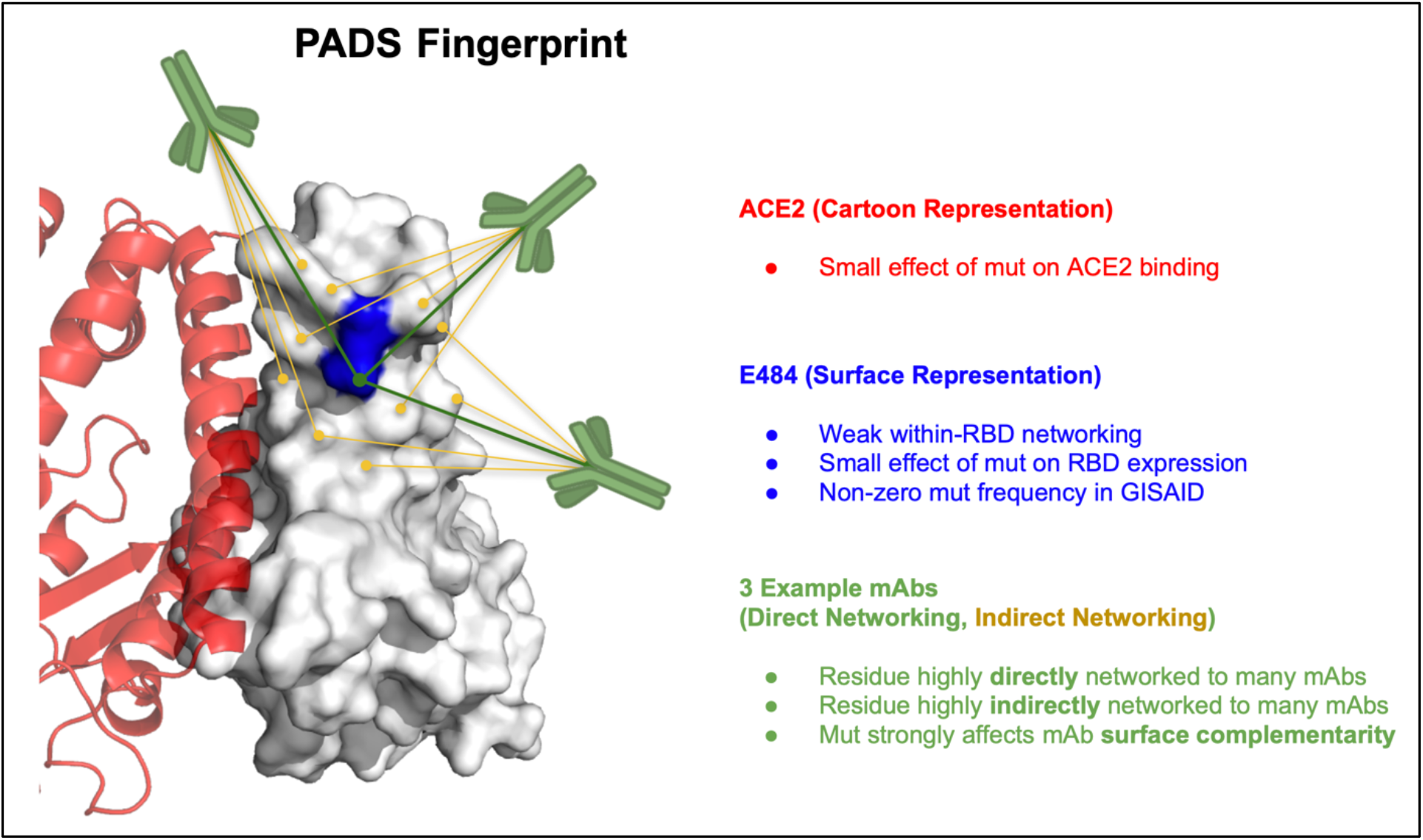
Canonical PADS Fingerprint. Mutation E484K is highlighted in blue on the RBD surface (PDB: 6M0J; [Lan et al. 2020]) and used to illustrate the fingerprint of a canonical RBD PAD. Three example mAbs are taken from the epitope-paratope map in Figure 1. Note that not all network connections nor the magnitude of the connections are shown for illustrative purposes. We find that antigenic and mutable residues tend to have a relatively small effect on fitness as measured by: ACE2 binding, ACE2 surface complementarity, RBD expression, within-RBD-networking, and mutation frequency in GISAID, and a relatively high antigenicity as measured by: direct networking, indirect networking, and perturbation of surface complementarity across the mAb/nanobody set.

**Table S1:**
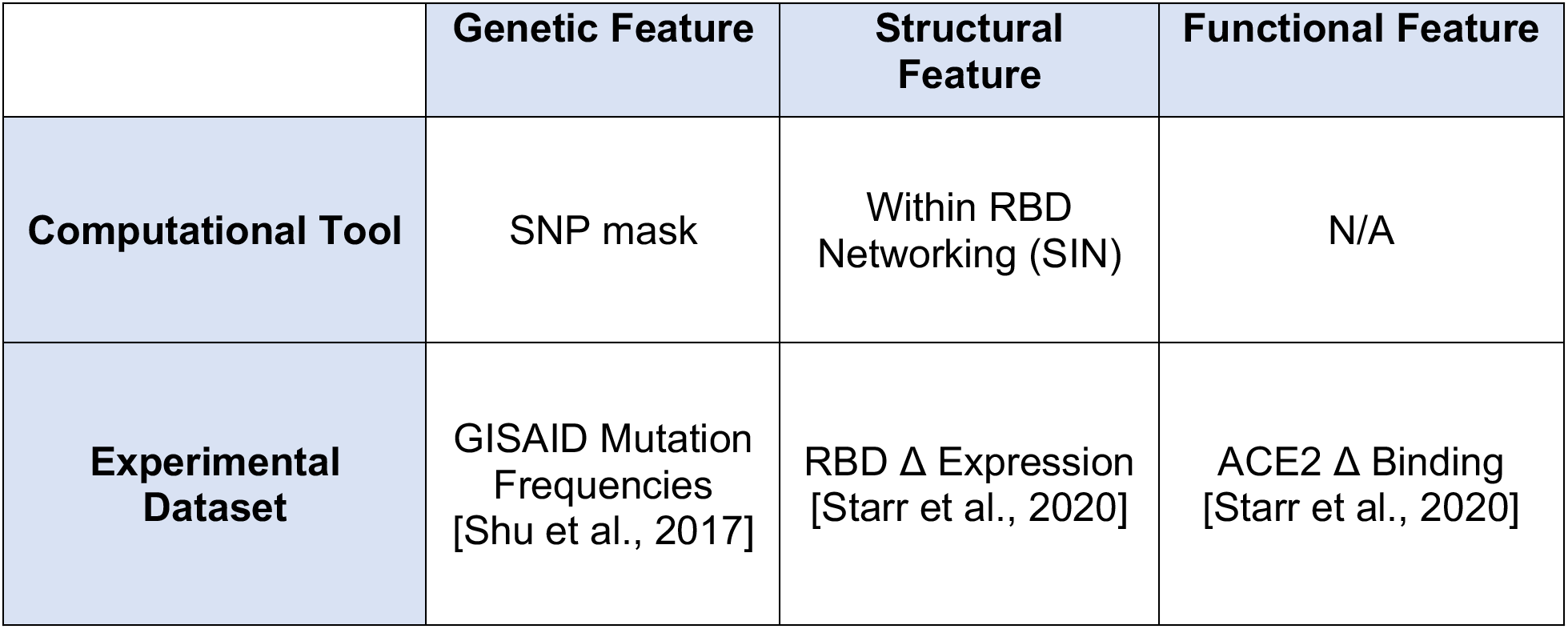
Features Used in Spectral Clustering of RBD Mutations Based on Mutability

**Table S2:**
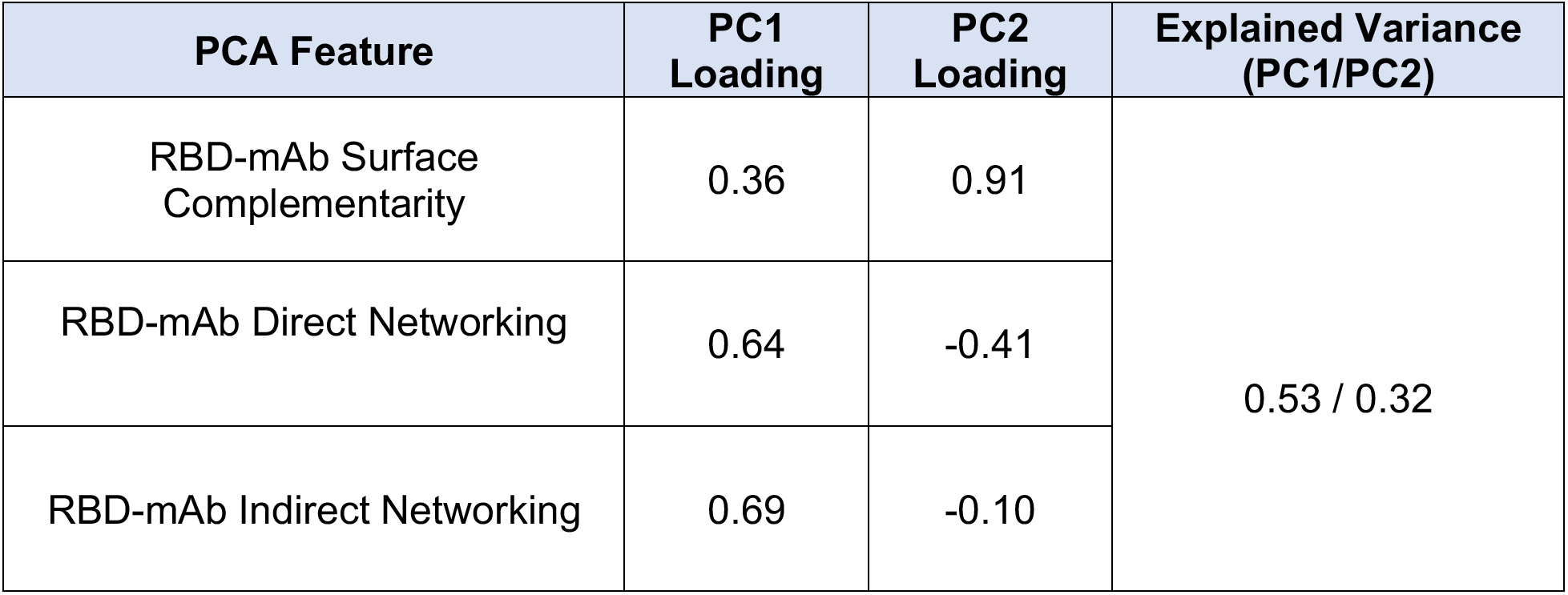
Antigenicity PCA Features and Fit

**Table S3:**
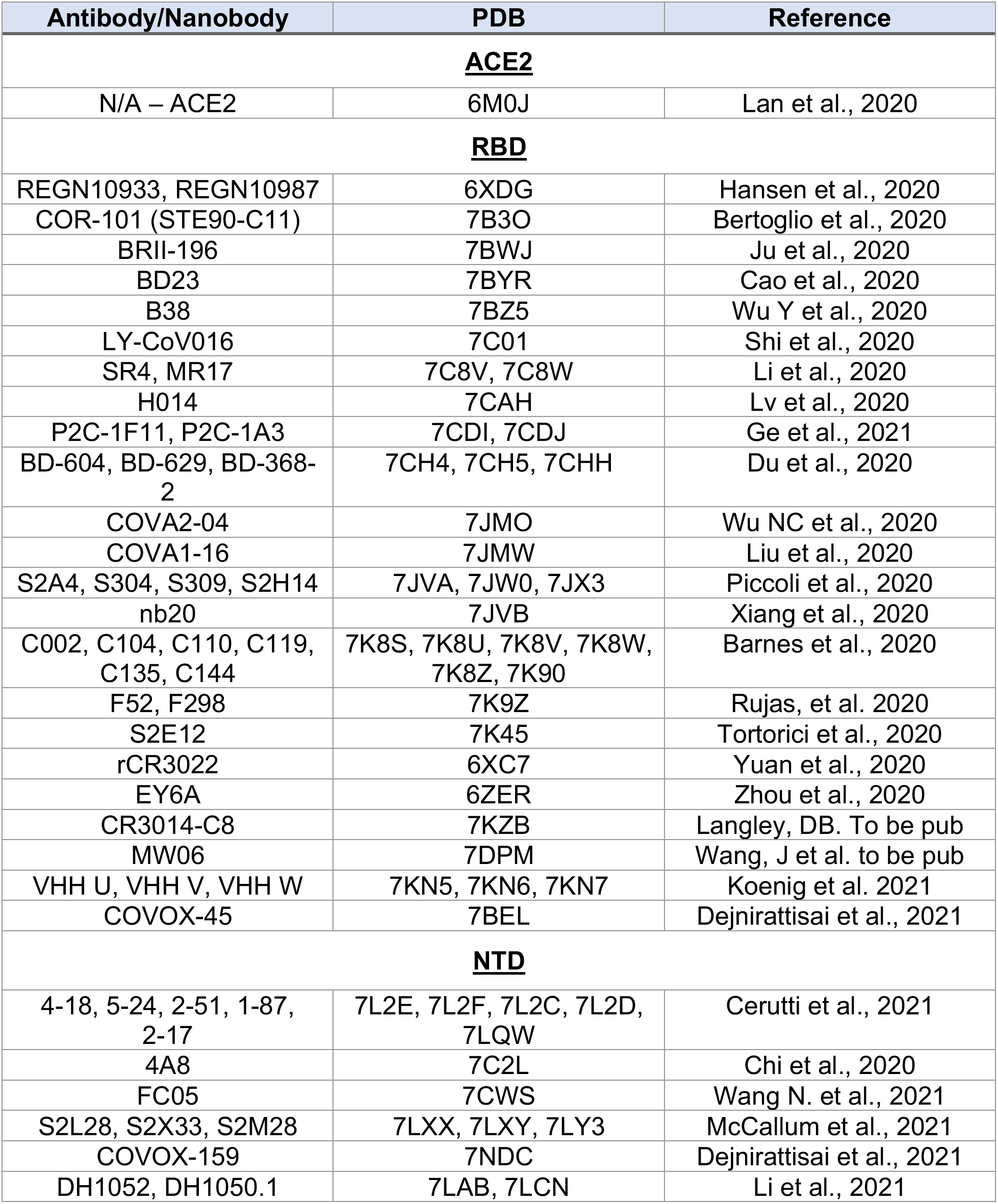
mAb/Nanobody – RBD/NTD Complex Structures

## Notes

### Competing Interest Statement

The authors have declared no competing interest.

## References

Abdool Karim SS, de Oliveira T. New SARS-CoV-2 Variants - Clinical, Public Health, and Vaccine Implications. N Engl J Med. 2021 May 13;384(19):1866–1868. doi: 10.1056/NEJMc2100362. Epub 2021 Mar 24. PMID: 33761203; PMCID: PMC8008749.

Abu-Raddad LJ, Chemaitelly H, Butt AA; National Study Group for COVID-19 Vaccination. Effectiveness of the BNT162b2 Covid-19 Vaccine against the B.1.1.7 and B.1.351 Variants. N Engl J Med. 2021 May 5:NEJMc2104974. doi: 10.1056/NEJMc2104974. Epub ahead of print. PMID: 33951357; PMCID: PMC8117967.

Barnes CO, Jette CA, Abernathy ME, Dam KA, Esswein SR, Gristick HB, Malyutin AG, Sharaf NG, Huey-Tubman KE, Lee YE, Robbiani DF, Nussenzweig MC, West AP Jr, Bjorkman PJ. SARS-CoV-2 neutralizing antibody structures inform therapeutic strategies. Nature. 2020 Dec;588(7839):682–687. doi: 10.1038/s41586-020-2852-1. Epub 2020 Oct 12. PMID: 33045718.

Bertoglio F, Fühner V, Ruschig M, Heine PA, Rand U,… Hust M et al. A SARS-CoV-2 neutralizing antibody selected from COVID-19 patients by phage display is binding to the ACE2-RBD interface and is tolerant to known RBD mutations. bioRxiv 2020.12.03.409318 (2020). https://doi.org/10.1101/2020.12.03.409318

Cathcart AL, Havenar-Daughton C, Lempp FA, Ma D, Schmid M, Agostini ML, Guarino B, Di iulio J, Rosen L, Tucker H, et al. The dual function monoclonal antibodies VIR-7831 and VIR-7832 demonstrate potent in vitro and in vivo activity against SARS-CoV-2. doi: https://doi.org/10.1101/2021.03.09.434607. Posted March 10, 2021.

Cao Y, Su B, Guo X, Sun W, Deng Y, Bao L, Zhu Q, Zhang X, Zheng Y, Geng C, et al. Potent Neutralizing Antibodies against SARS-CoV-2 Identified by High-Throughput Single-Cell Sequencing of Convalescent Patients’ B Cells. Cell. 2020 Jul 9;182(1):73–84.e16. doi: 10.1016/j.cell.2020.05.025. Epub 2020 May 18. PMID: 32425270; PMCID: PMC7231725.

Cerutti G, Guo Y, Zhou T, Gorman J, Lee M, Rapp M, … Shapiro L (2021). Potent SARS-CoV-2 neutralizing antibodies directed against spike N-terminal domain target a single supersite. Cell host & microbe, S1931-3128(21)00133–5. Advance online publication. https://doi.org/10.1016/j.chom.2021.03.005

Challen R, Brooks-Pollock E, Read JM, Dyson L, Tsaneva-Atanasova K, Danon L. Risk of mortality in patients infected with SARS-CoV-2 variant of concern 202012/1: matched cohort study. BMJ. 2021 Mar 9;372:n579. doi: 10.1136/bmj.n579. PMID: 33687922; PMCID: PMC7941603.

Chaudhury S, Lyskov S, Gray JJ. PyRosetta: a script-based interface for implementing molecular modeling algorithms using Rosetta, Bioinformatics, Volume 26, Issue 5, 1 March 2010, Pages 689–691, https://doi.org/10.1093/bioinformatics/btq007

Cherian S, Potdar V, Jadhav S, Yadav P, Gupta N, Das M, … Bhusan R. (2021). Convergent evolution of SARS-CoV-2 spike mutations, L452R, E484Q and P681R, in the second wave of COVID-19 in Maharashtra, India. bioRxiv 2021.04.22.440932. doi: https://doi.org/10.1101/2021.04.22.440932

Chi X, Yan R, Zhang J, Zhang G, Zhang Y, Hao M, Zhang Z, Fan P, Dong Y, Yang Y, Chen Z, Guo Y, Zhang J, Li Y, Song X, Chen Y, Xia L, Fu L, Hou L, Xu J, Yu C, Li J, Zhou Q, Chen W. A neutralizing human antibody binds to the N-terminal domain of the Spike protein of SARS-CoV-2. Science. 2020 Aug 7;369(6504):650–655. doi: 10.1126/science.abc6952. Epub 2020 Jun 22. PMID: 32571838; PMCID: PMC7319273.

Cobey S, Hensley SE. Immune history and influenza virus susceptibility. Curr Opin Virol. 2017 Feb;22:105–111. doi: 10.1016/j.coviro.2016.12.004. Epub 2017 Jan 12. PMID: 28088686; PMCID: PMC5467731.

Davies NG, Abbott S, Barnard RC, Jarvis CI, Kucharski AJ, Munday JD, Pearson CAB, Russell TW, Tully DC, Washburne AD, et al. Estimated transmissibility and impact of SARS-CoV-2 lineage B.1.1.7 in England. Science. 2021 Mar 3:eabg3055. doi: 10.1126/science.abg3055. Epub ahead of print. PMID: 33658326.

Dejnirattisai W, Zhou D, Ginn HM, Duyvesteyn HME, Supasa P, Case JB, Zhao Y, Walter TS, Mentzer AJ, Liu C, et al. The antigenic anatomy of SARS-CoV-2 receptor binding domain. Cell. 2021 Feb 18:S0092-8674(21)00221–X. doi: 10.1016/j.cell.2021.02.032. Epub ahead of print. PMID: 33756110; PMCID: PMC7891125.

Du S, Cao Y, Zhu Q, Yu P, Qi F, Wang G, Du X, Bao L, Deng W, Zhu H, et al. Structurally Resolved SARS-CoV-2 Antibody Shows High Efficacy in Severely Infected Hamsters and Provides a Potent Cocktail Pairing Strategy. Cell. 2020 Nov 12;183(4):1013–1023.e13. doi: 10.1016/j.cell.2020.09.035. Epub 2020 Sep 14. PMID: 32970990; PMCID: PMC7489885.

Eguia RT, Crawford KHD, Stevens-Ayers T, Kelnhofer-Millevolte L, Greninger AL, Englund JA, Boeckh MJ, Bloom JD. A human coronavirus evolves antigenically to escape antibody immunity. PLoS Pathog. 2021 Apr 8;17(4):e1009453. doi: 10.1371/journal.ppat.1009453. PMID: 33831132; PMCID: PMC8031418.

Ge J, Wang R, Ju B et al. Antibody neutralization of SARS-CoV-2 through ACE2 receptor mimicry. Nat Commun 12, 250 (2021). https://doi.org/10.1038/s41467-020-20501-9

Greaney AJ, Loes AN, Crawford KHD, Starr TN, Malone KD, Chu HY, Bloom JD. Comprehensive mapping of mutations in the SARS-CoV-2 receptor-binding domain that affect recognition by polyclonal human plasma antibodies. Cell Host Microbe. 2021a Mar 10;29(3):463–476.e6. doi: 10.1016/j.chom.2021.02.003. Epub 2021 Feb 8. PMID: 33592168; PMCID: PMC7869748.

Greaney AJ, Starr TN, Barnes CO, Weisblum Y, Schmidt F, Caskey M, Gaebler C, Cho A, Agudelo M, Finkin S, Wang Z, Poston D, Muecksch F, Hatziioannou T, Bieniasz PD, Robbiani DF, Nussenzweig MC, Bjorkman PJ, Bloom JD. Mutational escape from the polyclonal antibody response to SARS-CoV-2 infection is largely shaped by a single class of antibodies. bioRxiv [Preprint]. 2021b Mar 18:2021.03.17.435863. doi: 10.1101/2021.03.17.435863. PMID: 33758856; PMCID: PMC7987015.

Hansen J, Baum A, Pascal KE, Russo V, Giordano S, Wloga E, Fulton BO, Yan Y, Koon K, Patel Ket al. Studies in humanized mice and convalescent humans yield a SARS-CoV-2 antibody cocktail. Science. 2020 Aug 21;369(6506):1010–1014. doi: 10.1126/science.abd0827. Epub 2020 Jun 15. PMID: 32540901; PMCID: PMC7299284.

Hoffmann M, Arora P, Groß R, Seidel A, Hörnich BF, Hahn AS, Krüger N, Graichen L, Hofmann-Winkler H, Kempf A, Winkler MS, Schulz S, Jäck HM, Jahrsdörfer B, Schrezenmeier H, Müller M, Kleger A, Münch J, Pöhlmann S. SARS-CoV-2 variants B.1.351 and P.1 escape from neutralizing antibodies. Cell. 2021 Mar 20:S0092-8674(21)00367–6. doi: 10.1016/j.cell.2021.03.036. Epub ahead of print. PMID: 33794143; PMCID: PMC7980144.

Huang Y, Sun H, Yu H, Li S, Zheng Q, Xia N. Neutralizing antibodies against SARS-CoV-2: current understanding, challenge and perspective. Antib Ther. 2020;3(4):285–299. Published 2020 Dec 28. doi:10.1093/abt/tbaa028

Jiang S, Hillyer C, Du L. Neutralizing Antibodies against SARS-CoV-2 and Other Human Coronaviruses. Trends Immunol. 2020 May;41(5):355–359. doi: 10.1016/j.it.2020.03.007. Epub 2020 Apr 2. Erratum in: Trends Immunol. 2020 Apr 24;: PMID: 32249063; PMCID: PMC7129017.

Ju B, Zhang Q, Ge J, Wang R, Sun J, Ge X, Yu J, Shan S, Zhou B, Song S, et al. Human neutralizing antibodies elicited by SARS-CoV-2 infection. Nature. 2020 Aug;584(7819):115–119. doi: 10.1038/s41586-020-2380-z. Epub 2020 May 26. PMID: 32454513.

Kistler KE, Bedford T. Evidence for adaptive evolution in the receptor-binding domain of seasonal coronaviruses OC43 and 229e. Elife. 2021 Jan 19;10:e64509. doi: 10.7554/eLife.64509. PMID: 33463525; PMCID: PMC7861616.

Koenig PA, Das H, Liu H, Kümmerer BM, Gohr FN, Jenster LM, Schiffelers LDJ, Tesfamariam YM, Uchima M, Wuerth JD, et al. Structure-guided multivalent nanobodies block SARS-CoV-2 infection and suppress mutational escape. Science. 2021 Feb 12;371(6530):eabe6230. doi: 10.1126/science.abe6230. Epub 2021 Jan 12. PMID: 33436526.

Lan J, Ge J, Yu J, Shan S, Zhou H, Fan S, Zhang Q, Shi X, Wang Q, Zhang L, Wang X. Structure of the SARS-CoV-2 spike receptor-binding domain bound to the ACE2 receptor. Nature. 2020 May;581(7807):215–220. doi: 10.1038/s41586-020-2180-5. Epub 2020 Mar 30. PMID: 32225176.

Langley DB, Christ D. Potent SARS-CoV-2 binding and neutralization through maturation of iconic SARS-CoV-1 antibodies. To be published. https://www.rcsb.org/structure/7KZB

Li D, Edwards RJ, Manne K, Martinez DR, Schäfer A, Alam SM, Wiehe K, Lu X, Parks R, Sutherland LL, Oguin TH, McDanal C, Perez LG, Mansouri K, Gobeil SMC, Janowska K, Stalls V, Kopp M, Cai F, Lee E, Foulger A, Hernandez GE, Sanzone A, Tilahun K, Jiang C, Tse LV, Bock KW, Minai M, Nagata BM, Cronin K, Gee-Lai V, Deyton M, Barr M, Von Holle T, Macintyre AN, Stover E, Feldman J, Hauser BM, Caradonna TM, Scobey TD, Moody MA, Cain DW, DeMarco CT, Denny TN, Woods CW, Petzold EW, Schmidt AG, Teng IT, Zhou T, Kwong PD, Mascola JR, Graham BS, Moore IN, Seder R, Andersen H, Lewis MG, Montefiori DC, Sempowski GD, Baric RS, Acharya P, Haynes BF, Saunders KO. The functions of SARS-CoV-2 neutralizing and infection-enhancing antibodies in vitro and in mice and nonhuman primates. bioRxiv [Preprint]. 2021 Jan 2:2020.12.31.424729. doi: 10.1101/2020.12.31.424729. PMID: 33442694; PMCID: PMC7805451.

Li T, Cai H, Yao H, Zhou B, Zhang N, Gong Y, Zhao Y, Shen Q, Qin W, Hutter CAJ, et al. A potent synthetic nanobody targets RBD and protects mice from SARS-CoV-2 infection. bioRxiv; 2020. DOI: 10.1101/2020.06.09.143438.

Liu H, Wu NC, Yuan M, Bangaru S, Torres JL, Caniels TG, van Schooten J, Zhu X, Lee CD, Brouwer PJM, et al. Cross-Neutralization of a SARS-CoV-2 Antibody to a Functionally Conserved Site Is Mediated by Avidity. Immunity. 2020 Dec 15;53(6):1272–1280.e5. doi: 10.1016/j.immuni.2020.10.023. Epub 2020 Nov 25. PMID: 33242394; PMCID: PMC7687367.

Lv Z, Deng YQ, Ye Q, Cao L, Sun CY, Fan C, Huang W, Sun S, Sun Y, Zhu L, et al. Structural basis for neutralization of SARS-CoV-2 and SARS-CoV by a potent therapeutic antibody. Science. 2020 Sep 18;369(6510):1505–1509. doi: 10.1126/science.abc5881. Epub 2020 Jul 23. PMID: 32703908; PMCID: PMC7402622.

Madhi SA, Baillie V, Cutland CL, Voysey M, Koen AL, Fairlie L, Padayachee SD, Dheda K, Barnabas SL, Bhorat QE, et al. Efficacy of the ChAdOx1 nCoV-19 Covid-19 Vaccine against the B.1.351 Variant. N Engl J Med. 2021 Mar 16. doi: 10.1056/NEJMoa2102214. Epub ahead of print. PMID: 33725432.

McCallum M, Marco A, Lempp F, Tortorici MA, Pinto D, Walls AC, Beltramello M, Chen A, Liu Z, Zatta F, Zepeda S, di Iulio J, Bowen JE, Montiel-Ruiz M, Zhou J, Rosen LE, Bianchi S, Guarino B, Fregni CS, Abdelnabi R, Caroline Foo SY, Rothlauf PW, Bloyet LM, Benigni F, Cameroni E, Neyts J, Riva A, Snell G, Telenti A, Whelan SPJ, Virgin HW, Corti D, Pizzuto MS, Veesler D. N-terminal domain antigenic mapping reveals a site of vulnerability for SARS-CoV-2. bioRxiv [Preprint]. 2021 Jan 14:2021.01.14.426475. doi: 10.1101/2021.01.14.426475. Update in: Cell. 2021 Mar 16;: PMID: 33469588; PMCID: PMC7814825.

Pedregosa et al. Scikit-learn: Machine Learning in Python. JMLR 12, pp. 2825–2830, 2011.

Piccoli L, Park YJ, Tortorici MA, Czudnochowski N, Walls AC, Beltramello M, Silacci-Fregni C, Pinto D, Rosen LE, et al. Mapping Neutralizing and Immunodominant Sites on the SARS-CoV-2 Spike Receptor-Binding Domain by Structure-Guided High-Resolution Serology. Cell. 2020 Nov 12;183(4):1024–1042.e21. doi: 10.1016/j.cell.2020.09.037. Epub 2020 Sep 16. PMID: 32991844; PMCID: PMC7494283.

Pinto D, Park YJ, Beltramello M, Walls AC, Tortorici MA, Bianchi S, Jaconi S, Culap K, Zatta F, De Marco A, Peter A, Guarino B, Spreafico R, Cameroni E, Case JB, Chen RE, Havenar-Daughton C, Snell G, Telenti A, Virgin HW, Lanzavecchia A, Diamond MS, Fink K, Veesler D, Corti D. Cross-neutralization of SARS-CoV-2 by a human monoclonal SARS-CoV antibody. Nature. 2020 Jul;583(7815):290–295. doi: 10.1038/s41586-020-2349-y. Epub 2020 May 18. PMID: 32422645.

The PyMOL Molecular Graphics System, Version 2.0 Schrödinger, LLC. https://pymol.org/2/

Raybould MIJ, Kovaltsuk A, Marks C, Deane CM. CoV-AbDab: the Coronavirus Antibody Database [published online ahead of print, 2020 Aug 17]. Bioinformatics. 2020;btaa739. doi:10.1093/bioinformatics/btaa739

Robinson LN, Tharakaraman K, Rowley KJ, et al. Structure-Guided Design of an Anti-dengue Antibody Directed to a Non-immunodominant Epitope. Cell. 2015;162(3):493–504. doi:10.1016/j.cell.2015.06.057

Rees-Spear C, Muir L, Griffith SA, Heaney J, Aldon Y, Snitselaar JL, Thomas P, Graham C, Seow J, Lee N, et al. The effect of spike mutations on SARS-CoV-2 neutralization. Cell Rep. 2021 Mar 23;34(12):108890. doi: 10.1016/j.celrep.2021.108890. Epub 2021 Mar 6. PMID: 33713594; PMCID: PMC7936541.

Rujas E, Kucharska I, Tan YZ, Benlekbir S, Cui H, Zhao T, Wasney GA, Budylowski P, Guvenc F, Newton JC, et al. Multivalency transforms SARS-CoV-2 antibodies into broad and ultrapotent neutralizers. bioRxiv; 2020. DOI: 10.1101/2020.10.15.341636.

Sabino EC, Buss LF, Carvalho MPS, Prete CA Jr, Crispim MAE, Fraiji NA, Pereira RHM, Parag KV, da Silva Peixoto P, et al. Resurgence of COVID-19 in Manaus, Brazil, despite high seroprevalence. Lancet. 2021 Feb 6;397(10273):452–455. doi: 10.1016/S0140-6736(21)00183-5. Epub 2021 Jan 27. PMID: 33515491.

Shi R, Shan C, Duan X, Chen Z, Liu P, Song J, Song T, Bi X, Han C, Wu L, et al. A human neutralizing antibody targets the receptor-binding site of SARS-CoV-2. Nature. 2020 Aug;584(7819):120–124. doi: 10.1038/s41586-020-2381-y. Epub 2020 May 26. PMID: 32454512.

Shu Y, McCauley J. (2017) GISAID: Global initiative on sharing all influenza data – from vision to reality. EuroSurveillance, 22(13) DOI:10.2807/1560-7917.ES.2017.22.13.30494 PMCID: PMC5388101

Soundararajan V, Zheng S, Patel N, Warnock K, Raman R, Wilson IA, Raguram S, Sasisekharan V, Sasisekharan R. Networks link antigenic and receptor-binding sites of influenza hemagglutinin: mechanistic insight into fitter strain propagation. Sci Rep. 2011;1:200. doi: 10.1038/srep00200. Epub 2011 Dec 19. PMID: 22355715; PMCID: PMC3242012.

Starr TN, Greaney AJ, Hilton SK, Crawford KHD, Navarro MJ, Bowen JE, Tortorici MA, Walls AC, Veesler D, Bloom JD. Deep mutational scanning of SARS-CoV-2 receptor binding domain reveals constraints on folding and ACE2 binding. bioRxiv [Preprint]. 2020 Jun 17:2020.06.17.157982. doi: 10.1101/2020.06.17.157982. Update in: Cell. 2020 Aug 11;: PMID: 32587970; PMCID: PMC7310626.

Tortorici MA, Beltramello M, Lempp FA, Pinto D, Dang HV, Rosen LE, McCallum M, Bowen J, Minola A, Jaconi S, et al. Ultrapotent human antibodies protect against SARS-CoV-2 challenge via multiple mechanisms. Science. 2020 Nov 20;370(6519):950–957. doi: 10.1126/science.abe3354. Epub 2020 Sep 24. PMID: 32972994; PMCID: PMC7857395.

Voss WN, Hou YJ, Johnson NV, Delidakis G, Kim JE, Javanmardi K, Horton AP, Bartzoka F, Paresi CJ, Tanno Y, Chou CW, Abbasi SA, Pickens W, George K, Boutz DR, Towers DM, McDaniel JR, Billick D, Goike J, Rowe L, Batra D, Pohl J, Lee J, Gangappa S, Sambhara S, Gadush M, Wang N, Person MD, Iverson BL, Gollihar JD, Dye J, Herbert A, Finkelstein IJ, Baric RS, McLellan JS, Georgiou G, Lavinder JJ, Ippolito GC. Prevalent, protective, and convergent IgG recognition of SARS-CoV-2 non-RBD spike epitopes. Science. 2021 May 4:eabg5268. doi: 10.1126/science.abg5268. Epub ahead of print. PMID: 33947773.

Wang N, Sun Y, Feng R, Wang Y, Guo Y, Zhang L, Deng YQ, Wang L, Cui Z, Cao L, Zhang YJ, Li W, Zhu FC, Qin CF, Wang X. Structure-based development of human antibody cocktails against SARS-CoV-2. Cell Res. 2021 Jan;31(1):101–103. doi: 10.1038/s41422-020-00446-w. Epub 2020 Dec 1. PMID: 33262454; PMCID: PMC7705432.

Wang P, Liu L, Iketani S, Luo Y, Guo Y, Wang M, Yu J, Zhang B, Kwong PD, Graham BS, et al. Increased Resistance of SARS-CoV-2 Variants B.1.351 and B.1.1.7 to Antibody Neutralization. bioRxiv [Preprint]. 2021 Jan 26:2021.01.25.428137. doi: 10.1101/2021.01.25.428137. PMID: 33532778; PMCID: PMC7852271.

Wang J, Jiao S, Wang R, Zhang J, Zhang M, Wang M. 7DPM Crystal structure of SARS-CoV-2 Spike RBD in complex with MW06 Fab. Deposited: 2020-12-20 Released: 2021-02-17. To be published. DOI: 10.2210/pdb7DPM/pdb

Waskom M L, (2021). seaborn: statistical data visualization. Journal of Open Source Software, 6(60), 3021, https://doi.org/10.21105/joss.03021

Weisblum Y, Schmidt F, Zhang F, DaSilva J, Poston D, Lorenzi JC, Muecksch F, Rutkowska M, Hoffmann HH, Michailidis E, et al. Escape from neutralizing antibodies by SARS-CoV-2 spike protein variants. Elife. 2020 Oct 28;9:e61312. doi: 10.7554/eLife.61312. PMID: 33112236; PMCID: PMC7723407.

Wu Y, Wang F, Shen C, Peng W, Li D, Zhao C, Li Z, Li S, Bi Y, Yang Y, et al. A noncompeting pair of human neutralizing antibodies block COVID-19 virus binding to its receptor ACE2. Science. 2020 Jun 12;368(6496):1274–1278. doi: 10.1126/science.abc2241. Epub 2020 May 13. PMID: 32404477; PMCID: PMC7223722.

Wu NC, Yuan M, Liu H, Lee CD, Zhu X, Bangaru S, Torres JL, Caniels TG, Brouwer PJM, van Gils MJ, et al. An alternative binding mode of IGHV3-53 antibodies to the SARS-CoV-2 receptor binding domain. bioRxiv [Preprint]. 2020 Jul 27:2020.07.26.222232. doi: 10.1101/2020.07.26.222232. Update in: Cell Rep. 2020 Sep 29;:108274. PMID: 32743580; PMCID: PMC7386498.

Xiang Y, Nambulli S, Xiao Z, Liu H, Sang Z, Duprex WP, Schneidman-Duhovny D, Zhang C, Shi Y. Versatile and multivalent nanobodies efficiently neutralize SARS-CoV-2. Science. 2020 Dec 18;370(6523):1479-1484. doi: 10.1126/science.abe4747. Epub 2020 Nov 5. PMID: 33154108; PMCID: PMC7857400.

Yuan M, Liu H, Wu NC, Lee CD, Zhu X, Zhao F, Huang D, Yu W, Hua Y, Tien H, et al. Structural basis of a shared antibody response to SARS-CoV-2. Science. 2020 Aug 28;369(6507):1119–1123. doi: 10.1126/science.abd2321. Epub 2020 Jul 13. PMID: 32661058; PMCID: PMC7402627.

Zhou D, Duyvesteyn HME, Chen CP, Huang CG, Chen TH, Shih SR, Lin YC, Cheng CY, Cheng SH, Huang YC, et al. Structural basis for the neutralization of SARS-CoV-2 by an antibody from a convalescent patient. Nat Struct Mol Biol. 2020 Oct;27(10):950–958. doi: 10.1038/s41594-020-0480-y. Epub 2020 Jul 31. PMID: 32737466.

Zhou D, Dejnirattisai W, Supasa P, Liu C, Mentzer AJ, Ginn HM, Zhao Y, Duyvesteyn HME, Tuekprakhon A, Nutalai R, et al. Evidence of escape of SARS-CoV-2 variant B.1.351 from natural and vaccine-induced sera. Cell. 2021 Feb 23:S0092-8674(21)00226–9. doi: 10.1016/j.cell.2021.02.037. Epub ahead of print. PMID: 33730597; PMCID: PMC7901269.

Zhou D, Duyvesteyn HME, Chen CP, Huang CG, Chen TH, Shih SR, Lin YC, Cheng CY, Cheng SH, Huang YC, Lin TY, Ma C, Huo J, Carrique L, Malinauskas T, Ruza RR, Shah PNM, Tan TK, Rijal P, Donat RF, Godwin K, Buttigieg KR, Tree JA, Radecke J, Paterson NG, Supasa P, Mongkolsapaya J, Screaton GR, Carroll MW, Gilbert-Jaramillo J, Knight ML, James W, Owens RJ, Naismith JH, Townsend AR, Fry EE, Zhao Y, Ren J, Stuart DI, Huang KA. Structural basis for the neutralization of SARS-CoV-2 by an antibody from a convalescent patient. Nat Struct Mol Biol. 2020 Oct;27(10):950–958. doi: 10.1038/s41594-020-0480-y. Epub 2020 Jul 31. PMID: 32737466.

