## Supplementary Figures and Tables for "Mapping Potential Antigenic Drift Sites (PADS) on SARS-CoV-2 Spike in Continuous Epitope-Paratope Space"

### Supplemental Figures (S1-S7) and Tables (S1 to S3)

**Figure S1**: Correlation Between Indirect Networking of Mutable RBD Residues and RBD Sera Escape Mapping. Pearson correlation coefficient between sera escape [Greaney et al., 2021a] and indirect networking between mutable residues and paratopes of mAbs and nanobodies is $r=0.65$ ($p < .01$). For clarity, one outlying residue (E484) is removed from the bottom left graph, with approximate coordinates [0.19,0.14]. This point is shown on the top right graph, and is the highest scoring point for both sera escape and indirect networking.

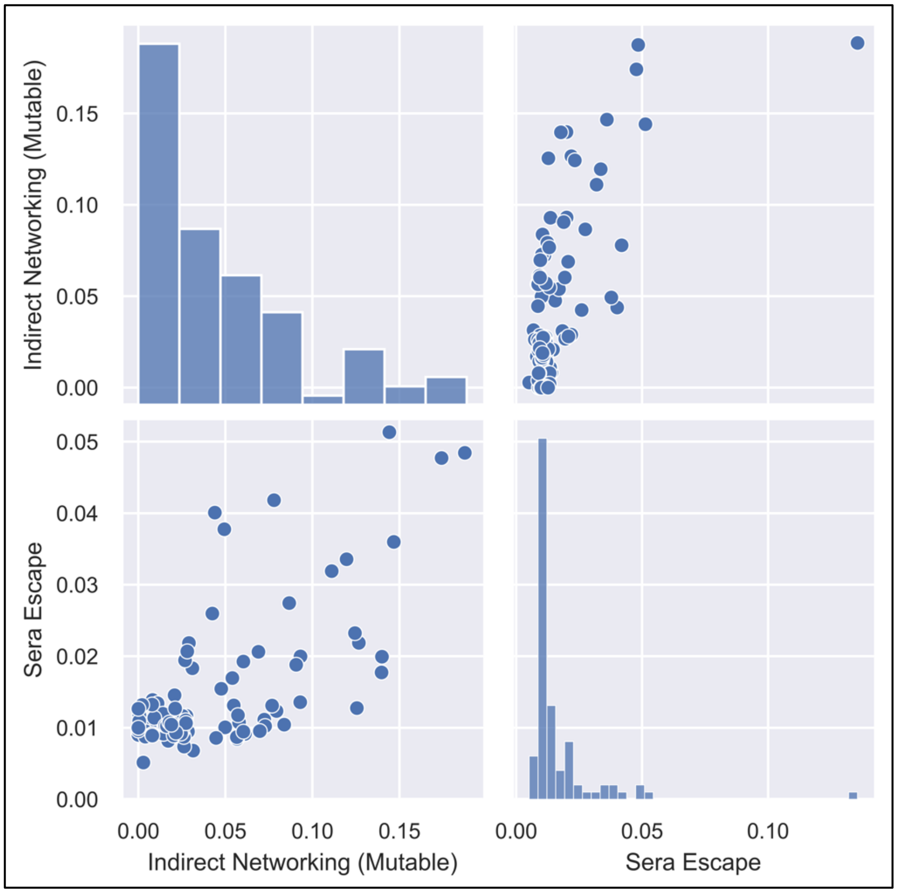

**Figure S2**: Total Epitope-Paratope Networking for N-Terminal Domain

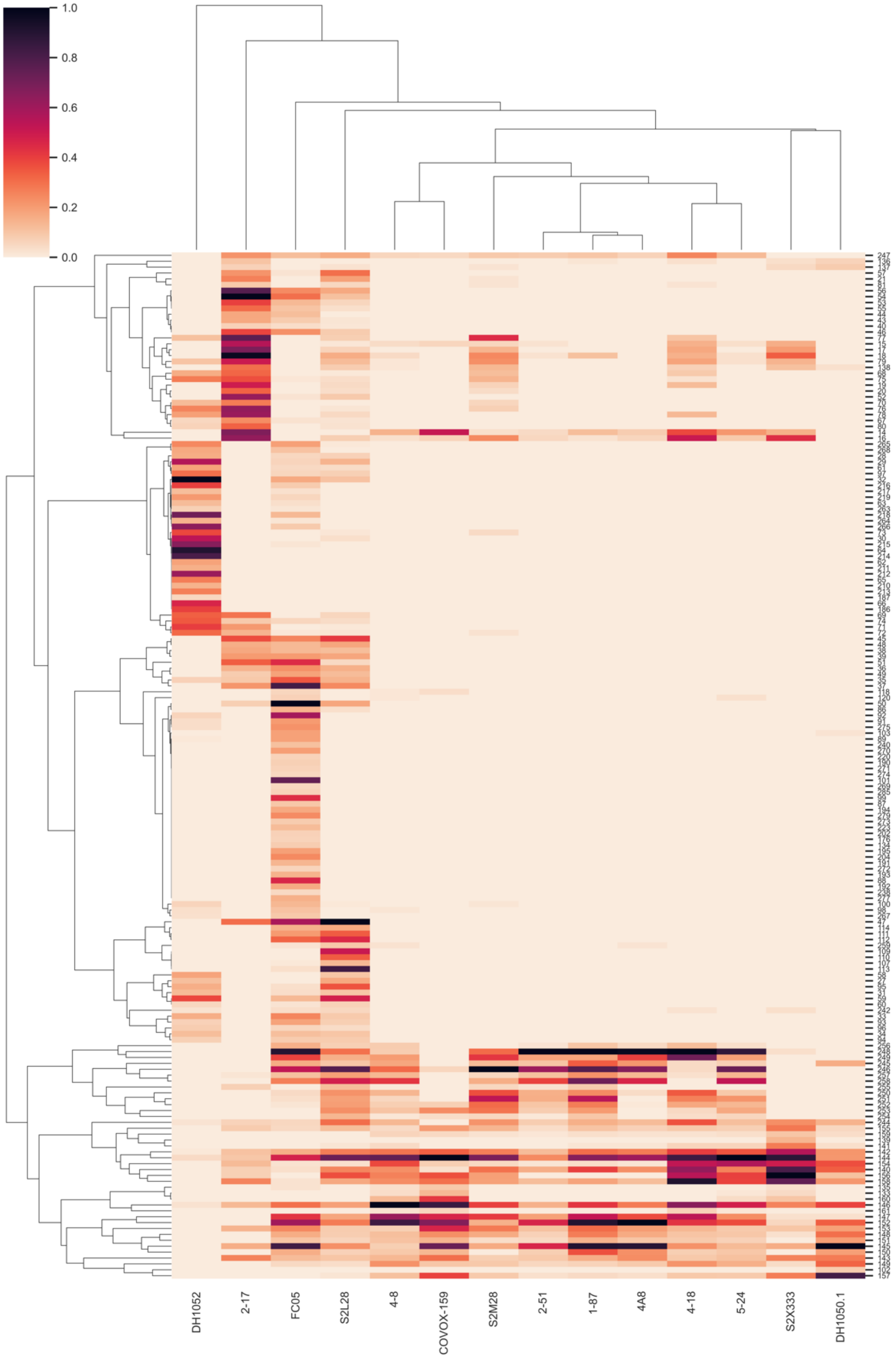

**Figure S3**: Direct Epitope-Paratope Networking for RBD

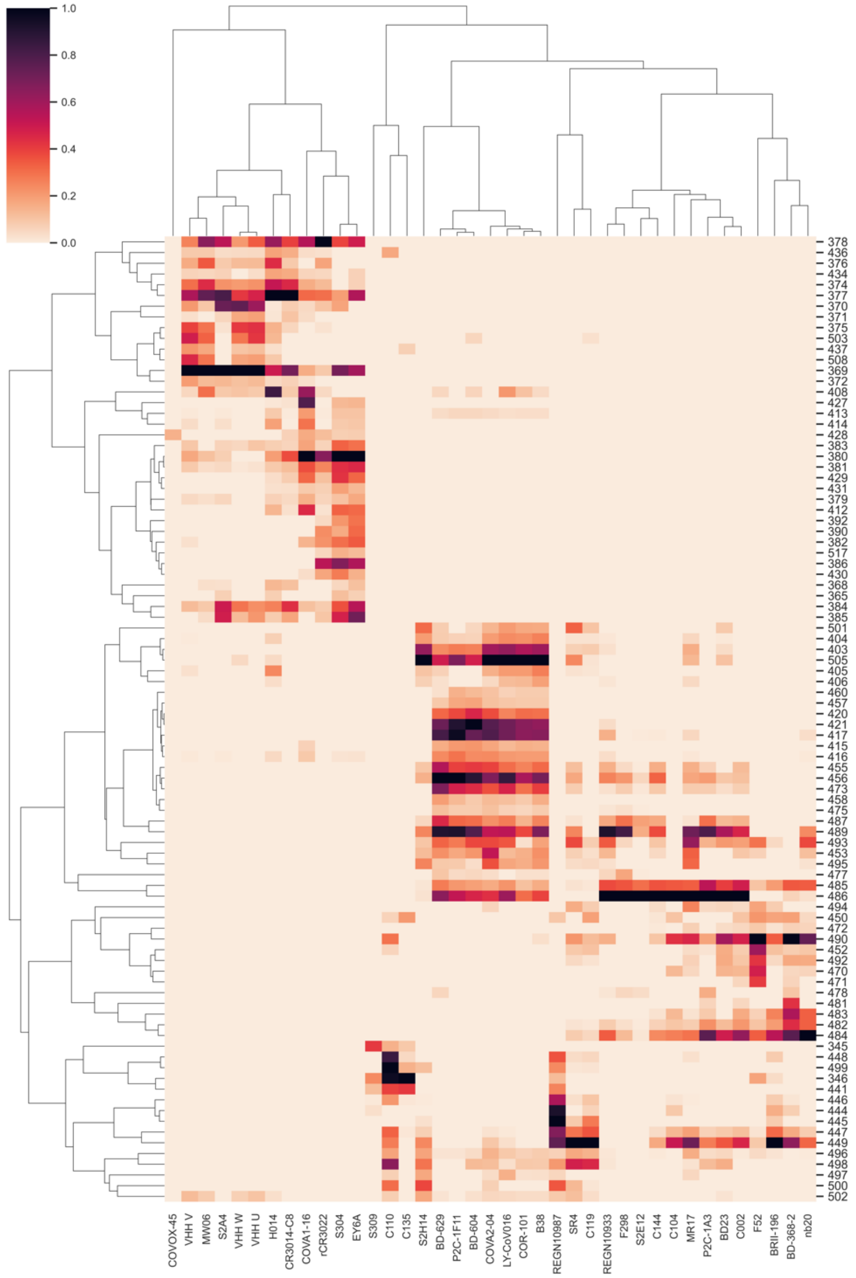

**Figure S4**: Indirect Epitope-Paratope Networking for RBD

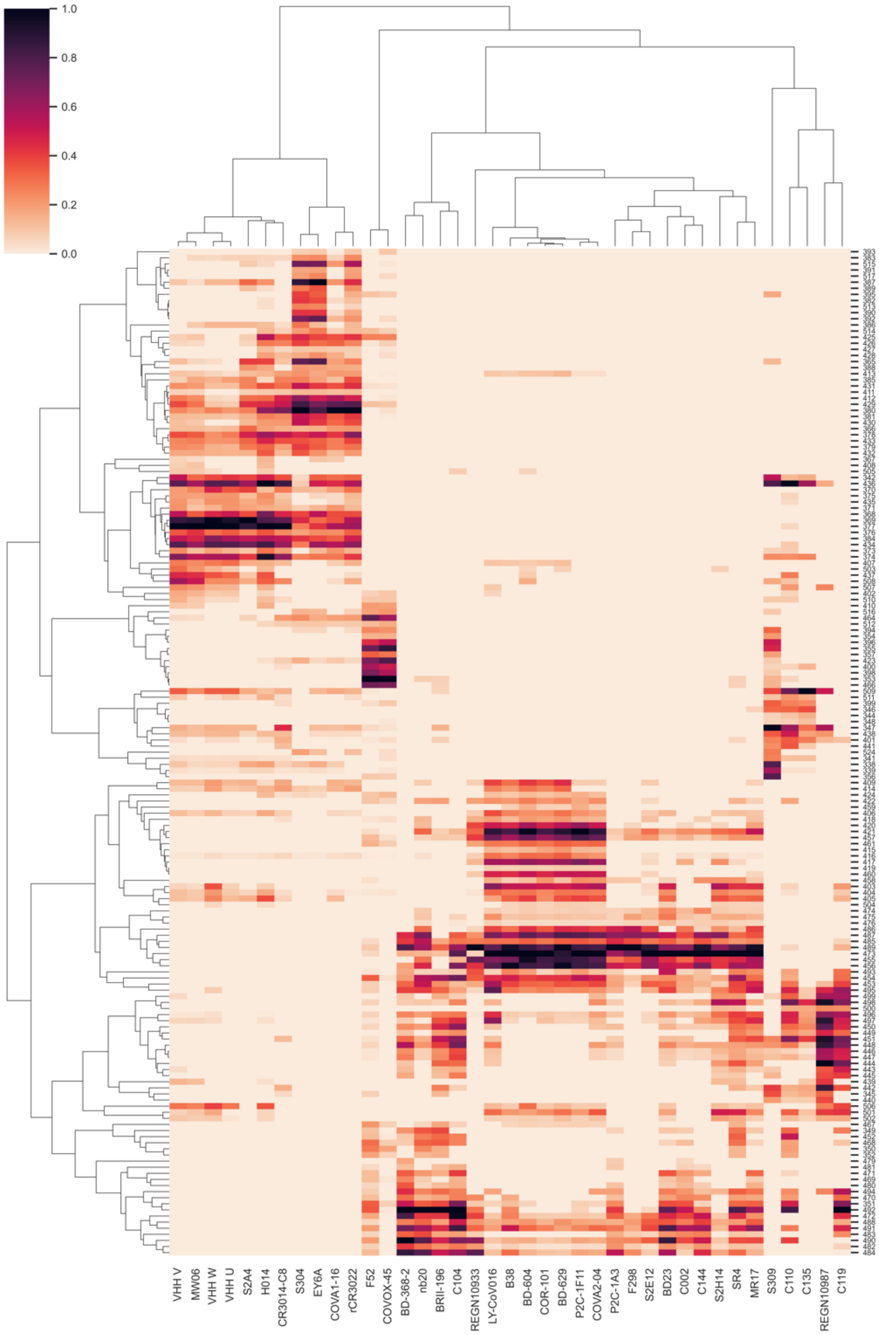

**Figure S5**: Mutability Clustering for all RBD SNP Mutations. Top, tSNE visualization of the clustering. Bottom, average and normalized (to absolute 1) values for each cluster along the feature set. Negative values imply knockdowns (e.g., in RBD expression, ACE2 binding, or ACE2 surface complementarity (SC)). Cluster 0 includes the RBD residues that appear the least genetically, structurally, and functionally constrained to mutate.

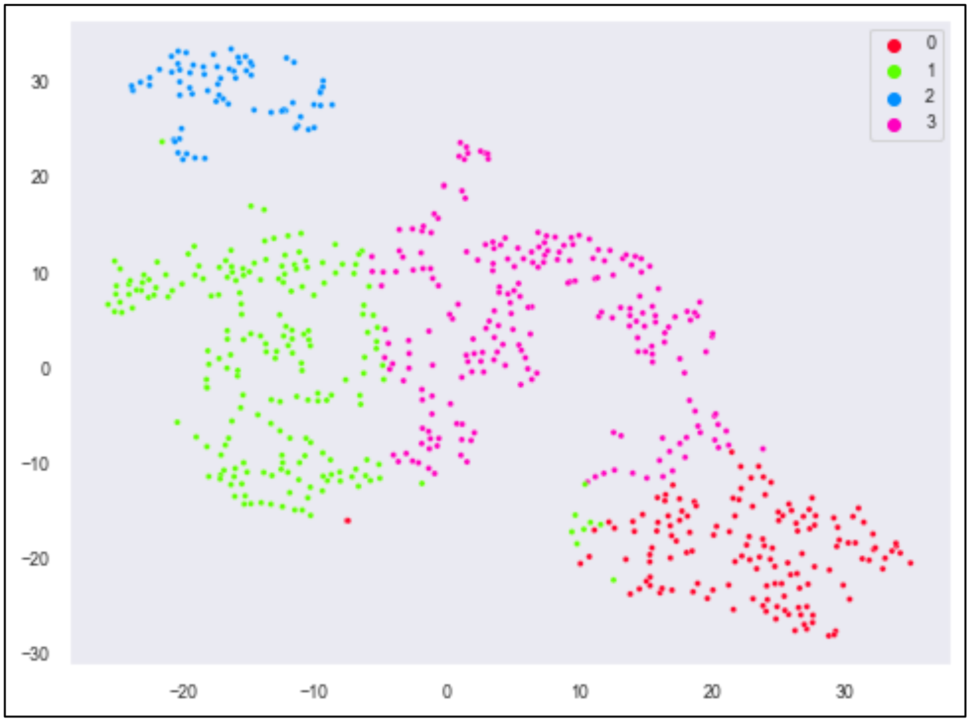


| Cluster | RBD Expression | ACE2 Binding | Within RBD Networking | GISAID % Observed |
|---|---|---|---|---|
| 0 | -0.15 | -0.03 | 0.20 | 100 |
| 1 | -0.52 | -0.09 | 0.35 | 14 |
| 2 | -0.26 | -0.84 | 0.28 | 7 |
| 3 | -0.06 | -0.04 | 0.17 | 17 |

**Figure S6:** Network model depicting the entire set of RBD PADS, instead of only the top PADS shown in Figure 2. See Figure 2 legend.

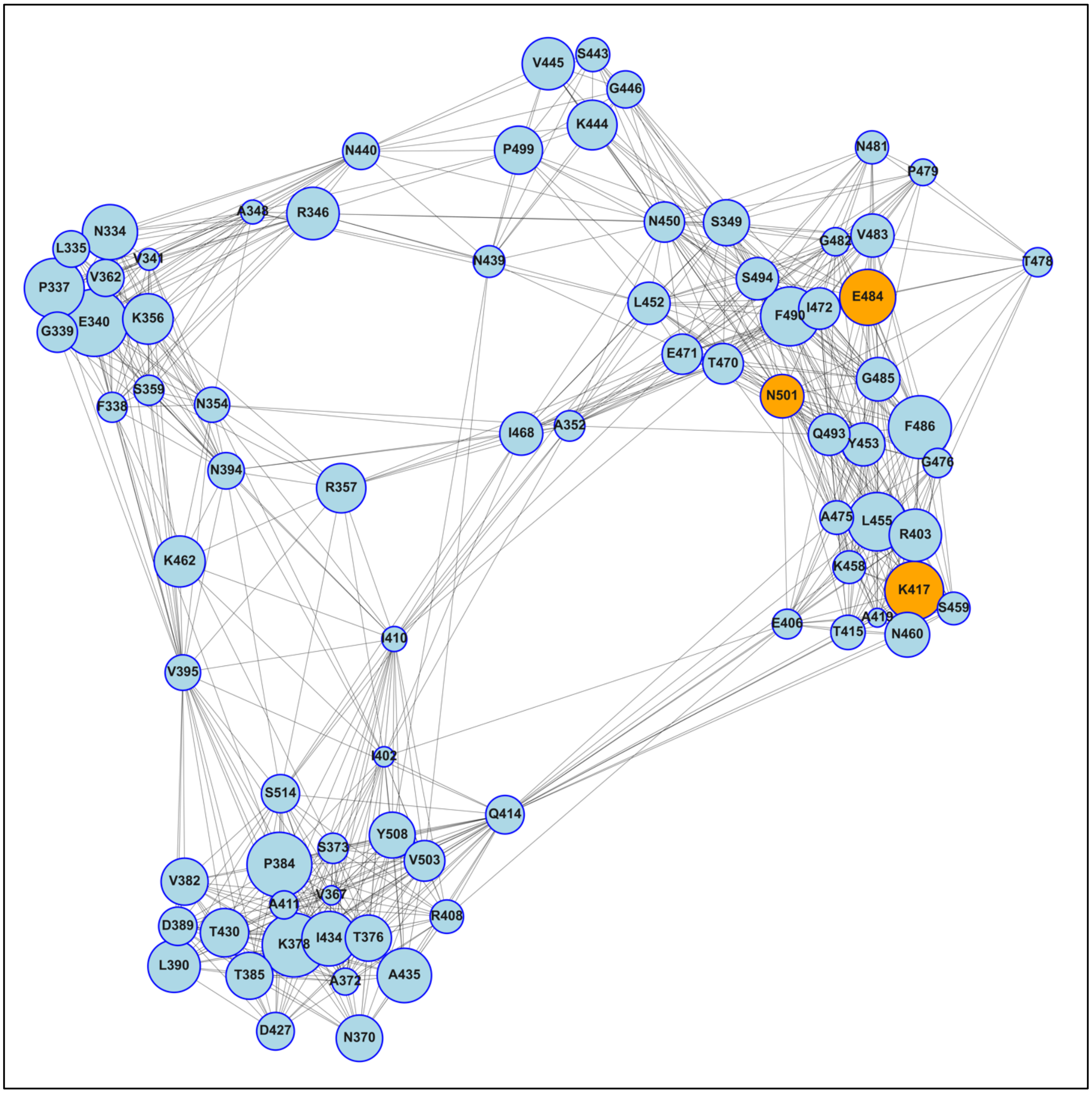

**Figure S7**: Canonical PADS Fingerprint. Mutation E484K is highlighted in blue on the RBD surface (PDB: 6M0J; [Lan et al. 2020]) and used to illustrate the fingerprint of a canonical RBD PAD. Three example mAbs are taken from the epitope-paratope map in Figure 1. Note that not all network connections nor the magnitude of the connections are shown for illustrative purposes. We find that antigenic and mutable residues tend to have a relatively small effect on fitness as measured by: ACE2 binding, ACE2 surface complementarity, RBD expression, within-RBD-networking, and mutation frequency in GISAID, and a relatively high antigenicity as measured by: direct networking, indirect networking, and perturbation of surface complementarity across the mAb/nanobody set.

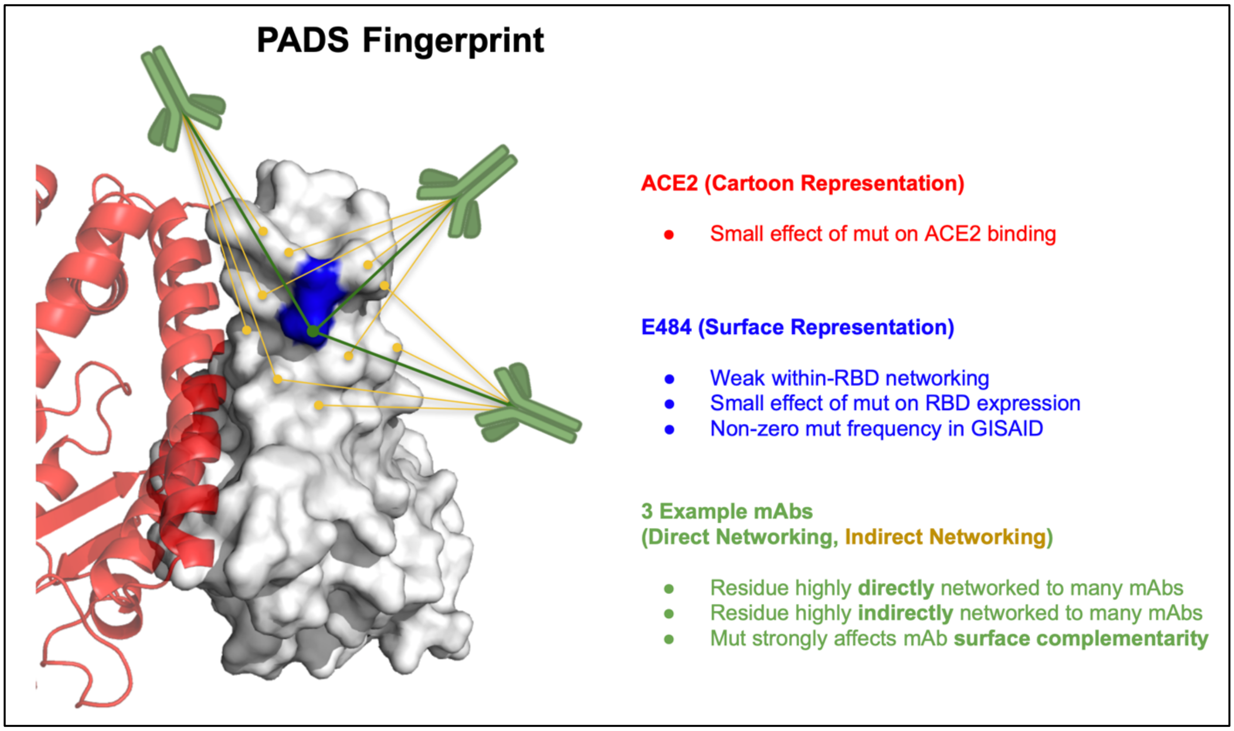

**Table S1**: Features Used in Spectral Clustering of RBD Mutations Based on Mutability

| | Genetic Feature | Structural Feature | Functional Feature |
|---|---|---|---|
| **Computational Tool** | SNP mask | Within RBD Networking (SIN) | N/A |
| **Experimental Dataset** | GISAID Mutation Frequencies [Shu et al., 2017] | RBD Δ Expression [Starr et al., 2020] | ACE2 Δ Binding [Starr et al., 2020] |

**Table S2**: Antigenicity PCA Features and Fit

| PCA Feature | PC1 Loading | PC2 Loading | Explained Variance (PC1/PC2) |
|---|---|---|---|
| RBD-mAb Surface Complementarity | 0.36 | 0.91 | 0.53 / 0.32 |
| RBD-mAb Direct Networking | 0.64 | -0.41 | |
| RBD-mAb Indirect Networking | 0.69 | -0.10 | |

**Table S3**: mAb/Nanobody – RBD/NTD Complex Structures

| Antibody/Nanobody | PDB | Reference |
|---|---|---|
| | **ACE2** | |
| N/A – ACE2 | 6M0J | Lan et al., 2020 |
| | **RBD** | |
| REGN10933, REGN10987 | 6XDG | Hansen et al., 2020 |
| COR-101 (STE90-C11) | 7B3O | Bertoglio et al., 2020 |
| BRII-196 | 7BWJ | Ju et al., 2020 |
| BD23 | 7BYR | Cao et al., 2020 |
| B38 | 7BZ5 | Wu Y et al., 2020 |
| LY-CoV016 | 7C01 | Shi et al., 2020 |
| SR4, MR17 | 7C8V, 7C8W | Li et al., 2020 |
| H014 | 7CAH | Lv et al., 2020 |
| P2C-1F11, P2C-1A3 | 7CDI, 7CDJ | Ge et al., 2021 |
| BD-604, BD-629, BD-368-2 | 7CH4, 7CH5, 7CHH | Du et al., 2020 |
| COVA2-04 | 7JMO | Wu NC et al., 2020 |
| COVA1-16 | 7JMW | Liu et al., 2020 |
| S2A4, S304, S309, S2H14 | 7JVA, 7JW0, 7JX3 | Piccoli et al., 2020 |
| nb20 | 7JVB | Xiang et al., 2020 |
| C002, C104, C110, C119, C135, C144 | 7K8S, 7K8U, 7K8V, 7K8W, 7K8Z, 7K90 | Barnes et al., 2020 |
| F52, F298 | 7K9Z | Rujas, et al. 2020 |
| S2E12 | 7K45 | Tortorici et al., 2020 |
| rCR3022 | 6XC7 | Yuan et al., 2020 |
| EY6A | 6ZER | Zhou et al., 2020 |
| CR3014-C8 | 7KZB | Langley, DB. To be pub |
| MW06 | 7DPM | Wang, J et al. to be pub |
| VHH U, VHH V, VHH W | 7KN5, 7KN6, 7KN7 | Koenig et al. 2021 |
| COVOX-45 | 7BEL | Dejnirattisai et al., 2021 |
| | **NTD** | |
| 4-18, 5-24, 2-51, 1-87, 2-17 | 7L2E, 7L2F, 7L2C, 7L2D, 7LQW | Cerutti et al., 2021 |
| 4A8 | 7C2L | Chi et al., 2020 |
| FC05 | 7CWS | Wang N. et al., 2021 |
| S2L28, S2X33, S2M28 | 7LXX, 7LXY, 7LY3 | McCallum et al., 2021 |
| COVOX-159 | 7NDC | Dejnirattisai et al., 2021 |
| DH1052, DH1050.1 | 7LAB, 7LCN | Li et al., 2021 |
